# The unfolded protein response links tumor aneuploidy to local immune dysregulation

**DOI:** 10.1101/2020.11.10.377358

**Authors:** Su Xian, Stephen Searles, Paras Sahani, T. Cameron Waller, Kristen Jepsen, Hannah Carter, Maurizio Zanetti

**Affiliations:** The Laboratory of Immunology, Department of Medicine and Moores Cancer Center, University of California San Diego, La Jolla, CA 92093 (USA); Division of Medical Genetics Biostatistics, Department of Medicine, Bioinformatics and System Biology Program, University of California, San Diego, La Jolla, CA 92093 (USA); IGM Genomics Center, University of California, San Diego, La Jolla, CA 92093 (USA)

**Author notes:** **Correspondence to**: Maurizio Zanetti, Hannah Carter.

## Abstract

Aneuploidy is a chromosomal abnormality associated with poor prognosis in many cancer types. Here we tested the hypothesis that the unfolded protein response (UPR) links mechanistically aneuploidy and local immune dysregulation. Using a single somatic copy-number alteration (SCNA) score inclusive of whole-chromosome, arm and focal chromosome alterations in a pan-cancer analysis of 9,375 samples in The Cancer Genome Atlas (TCGA) database, we found an inverse correlation with a cytotoxicity (CYT) score across disease stages. Co-expression patterns of UPR genes changed substantially between SCNA^low^ and SCNA^high^ groups. Pathway activity scores showed increased activity by multiple branches of the UPR in response to aneuploidy. The PERK branch showed the strongest association with a reduced CYT score. The conditioned medium of aneuploid cells transmitted XBP1 splicing and caused IL-6 and Arginase1 transcription in receiver bone marrow-derived macrophages. We propose the UPR as a mechanistic link between aneuploidy and immune dysregulation in the tumor microenvironment.

**Statement of Significance:** Aneuploidy accumulates over the life of a tumor and is associated with poor prognosis. Tumor progression is also associated with a progressive immune dysregulation. To explain these complex and concurrent disorders we tested the hypothesis that the unfolded protein could represent the link between aneuploidy and a dysregulation of local immunity favoring tumor progression.

## Introduction

Aneuploidy is the oldest form of chromosomal abnormality identified (Boveri, 2008) and can result from mis-segregation during anaphase (e.g., spindle assembly, checkpoint defects) (Gordon et al., 2012), cell fusion (Migeon et al., 1974) or cell-in-cell formation (entosis) (Krajcovic et al., 2011). In cancer aneuploidy is part of a broader category of genomic abnormalities called somatic copy-number alteration (SCNA; distinguished from germline copy-number variations). Approximately 90% of solid tumors and 50% of blood cancers present some features of aneuploidy (Beroukhim et al., 2010; Mitelman et al., 2016). SCNA can be divided into three categories: whole-chromosome, chromosome-arm and focal (Beroukhim et al., 2010). Aneuploidy is associated with tumor progression and poor prognosis (Hieronymus et al., 2018; Newburger et al., 2013; Owainati et al., 1987; Stopsack et al., 2019) and chromosomally unstable cancer cells exhibit increased multidrug resistance (Duesberg et al., 2000). Aneuploidy, which is usually detrimental to cell viability in healthy tissues leading to negative selection of aneuploid cells, is paradoxically tolerated in cancer cells (Holland and Cleveland, 2009; Valind et al., 2013; Varetti et al., 2014) suggesting that it provides selective growth advantage to cancer cells in the hostile metabolic tumor microenvironment (Giam and Rancati, 2015).

Recent reports showed that tumor aneuploidy correlates with markers of immune evasion (Davoli et al., 2017) and reduced number of tumor-infiltrating leukocytes (Taylor et al., 2018) suggesting a connection between aneuploidy and immune surveillance. However, neither study provided a mechanistic explanation for the correlation. Paradoxically, two earlier reports (Boileve et al., 2013; Senovilla et al., 2012) showed that tetraploid neoplastic cells ostensibly lead to their selective elimination by T cells. A negative impact of aneuploidy on local immunity represents therefore an important new variable in the interplay between cancer and immunity.

Aneuploidy is a source of genetic variation allowing for evolutionarily selection and advantage (Torres et al., 2007) but this may also have a functional impact on cells through gene and protein dosage change as demonstrated in yeast and in mammalian cells (Pavelka et al., 2010; Sheltzer et al., 2012; Stranger et al., 2007). Furthermore, in yeast, quantitative changes in the proteome beyond the buffering capability of the cell were shown to cause an unfolded protein response (UPR) (Geiler-Samerotte et al., 2011) and hypo-osmotic stress (Tsai et al., 2019). Congruently, a proteotoxic response was documented in human aneuploid non-cancer cells (Stingele et al., 2012) and predicted to be a consequence of aneuploidy in cancer cells (Chunduri and Storchova, 2019; Zhu et al., 2018).

The UPR is mediated by three initiator/sensor ER transmembrane molecules: inositol-requiring enzyme 1 (IRE1α), PKR-like ER kinase (PERK), and activating transcription factor 6 (ATF6), which are maintained inactive through association with the 78-kDa glucose-regulated protein (GRP78) (Schroder and Kaufman, 2005). During ER stress, GRP78 disassociates from each of the three sensor molecules, activating downstream signaling cascades to normalize protein folding and secretion. PERK phosphorylates translation initiation factor 2 (eIF2α), resulting in global inhibition of translation to reduce ER client proteins (Walter and Ron, 2011). IRE1α auto-phosphorylates to activate its endonuclease domain, resulting in the cleavage of *XBP1* to generate a spliced XBP1 isoform (XBP1s), which drives the production of various ER chaperones to restore ER homeostasis (Walter and Ron, 2011). XBP1s also binds to the promoter of several pro-inflammatory cytokine genes (Martinon et al., 2010). In addition, under ER stress or forced autophosphorylation, IRE1α’s RNase domain can cause endonucleolytic decay of many ER-localized mRNAs through a phenomenon termed regulated IRE1-dependent decay (RIDD) (Hollien and Weissman, 2006). ATF6 induces XBP1 and translocates to the Golgi where it is cleaved into its functional form, and acts in parallel with XBP1s to restore ER homeostasis as a transcription factor (Yoshida et al., 2001). If these compensatory mechanisms fail, downstream signaling from PERK via transcription factor 4 (ATF4) activates the transcription factor CCAAT-enhancer-binding protein homologous protein (CHOP) encoded by *DDIT3* to initiate apoptosis (Walter and Ron, 2011).

In cancer cells the UPR serves as a cell-autonomous process to restore proteostasis, enable survival and signal cell growth (Clarke et al., 2014; Lee, 2014). However, it can also function cell-nonautonomously by promoting the release of soluble molecules that target neighboring cells (Mahadevan et al., 2011; Rodvold et al., 2017). These can increase the fitness and survival of tumor cells (Rodvold et al., 2017), impart immunosuppressive and pro-tumorigenic functions to bone marrow-derived macrophages and dendritic cells (Cubillos-Ruiz et al., 2015; Mahadevan et al., 2012; Mahadevan et al., 2011), and impair indirectly the function of T cells (Mahadevan et al., 2012; Song et al., 2018). Because the link between aneuploidy, UPR and immune cells, has not been interrogated, here we tested the hypothesis that the UPR may represent the link between aneuploidy and immune dysregulation in the tumor microenvironment (Zanetti, 2017). To this end, we applied statistical methods to UPR gene expression and pathway structures in a pan-cancer analysis of 9,375 TCGA samples across 32 tumor types using a SCNA score (inclusive of whole-chromosome, arm and focal SCNA), and analyzed the effects of aneuploidy generated *in vitro* on the UPR and the induction of a mixed proinflammatory/immune suppressive phenotype in bone marrow derived macrophages. We show that the UPR is a mechanism by which aneuploidy may disrupt local immunity in cancer contributing to the loss of immune surveillance.

## Results

### Pan-cancer distribution of SCNA

SCNA has been previously grouped into three categories: whole-chromosome, arm and focal (Beroukhim et al., 2010; Davoli et al., 2017). Here we used a similar distinction (Fig. 1A) where <u>whole-chromosome</u> copy-number alteration refers to a duplication or loss of an entire chromosome (canonical aneuploidy), <u>arm</u> copy-number alteration refers to the duplication or loss of an entire chromosome arm, and <u>focal</u> copy-number alteration refers to the duplication or loss of a discrete region of the chromosome not spanning the length of the entire chromosome arm. In a typical cancer sample, arm and focal SCNA have been estimated at 25% and 10% of the genome, respectively (Beroukhim et al., 2010; Gordon et al., 2012). Here we studied the 3 types of SCNA in TCGA, quantifying them from segmented SNP array intensity data (see Methods). An example of a segmented SNP profile of a single tumor harboring all three event categories is shown in Figure 1B.

**Figure 1.**
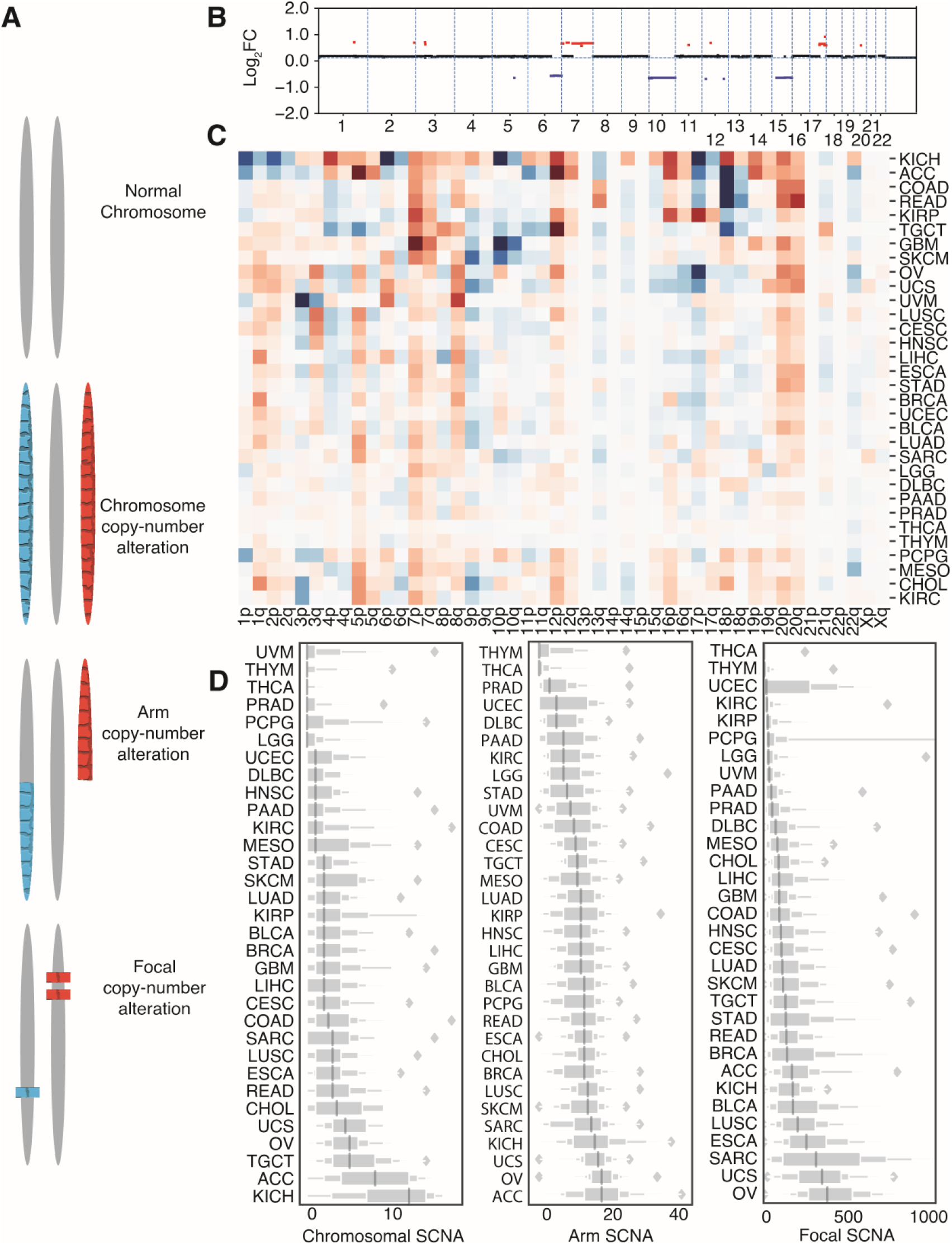
Measuring tumor aneuploidy. (**A**) An illustration on the definition of focal, arm, and chromosome level somatic copy-number alterations (SCNAs). (**B**) Example of the somatic copy-number alteration distribution in a single TCGA sample (TCGA-02-0003, GBM) with red suggesting copy number gain and blue suggesting copy number loss. The x axis represents the chromosomal location and the y axis stands for log_2_ fold change value of the corresponding region. (**C**) Heatmap showing patterns of copy number gain (red) or loss (blue) of chromosomal p and q arms across 32 tumor types in TCGA. (**D**) Box plots showing total counts of copy number alteration events (either loss or gain) across 32 tumor types in TCGA. From left to right, plots depict events of focal, arm or chromosome

We first studied the distribution of SCNAs across thirty-two different tumor types (n = 9,375) from TCGA. In a heatmap depicting only arm level copy-number alterations, we observed different patterns of alteration for different tumor types (Fig. 1C). Both 3p arm losses and 3q arm gains were evident in lung squamous cell carcinoma (LUSC) (Fig. 1C), consistent with a previous report (Zabarovsky et al., 2002). We further compared the frequencies by tumor of the three categories of SCNA across the thirty-two tumor types (Fig. 1D). Some tumor types such as thyroid carcinoma (THCA) and thymoma (THYM), show low frequency of SCNA, while others such as ovarian serous adenocarcinoma (OV) and kidney chromophobe cancer (KICH) carry heavy SCNA burdens (Fig. 1D). Previous studies suggested an inverse correlation between the number of non-synonymous mutations and copy number alterations (Ciriello et al., 2013), yet another study suggested that this inverse correlation might derive primarily from the microsatellite instability high (MSI-H) group (Taylor et al., 2018). Our data agree with this latter interpretation (Fig. S1). Moreover, we found a positive association between *TP53* mutations and SCNA scores (Fig. S2A), consistent with the role of *TP53* in protecting against chromosome segregation errors (Soto et al., 2017). We computed *TP53* activity scores using ten *TP53* repressed genes (Cancer Genome Atlas Research Network. Electronic address and Cancer Genome Atlas Research, 2017) and found a significant negative correlation between *TP53* activity and SCNA scores in seventeen out of thirty-two cancer types (Fig. S2B). Among these, only THYM showed a significant positive correlation. Our results support the view that inactivation and mutations of *TP53* associate with high SCNA scores in most solid tumors (Zack et al., 2013).

### A single SCNA score negatively correlates with immune-mediated cytotoxicity

We first sought to determine if the three types of SCNA could be used and expressed collectively as a single SCNA score. We used pairwise correlation to evaluate the relationship between whole-chromosome, arm and focal SCNA categories (raw event count; Spearman correlation, Fig. S3A) and found a strong, positive inter-category correlation (Spearman r= 0.548-0.627). We then derived aggregate scores for each category separately and compared them to a single combined SCNA score (see also Methods). The combined SCNA score showed consistently high correlation with all three categories considered independently (Spearman r= 0.735-0.866) with focal SCNA being the least correlated (Spearman r = 0.735) (Fig. S3A).

In a pan-cancer analysis of tumors with stage information (n = 6298, 25 tumor types), we found that as tumor stage increased, the single combined SCNA score (aneuploidy score) also increased (Fig. 2A). We also measured perforin (*PRF1*) and granzyme A (*GZMA*) gene expression as representations of cytolytic activity (CYT) in tumors (Rooney et al., 2015) and found that CYT was inversely correlated with tumor stages across all cancer types (n =6458, 25 tumor types) (Fig. 2B). To account for a potential bias due to differences in stage and SCNA distribution across tumor types (Fig. 1D) we included tumor type as a covariate in an Ordinary Least Squares (OLS) linear regression model. We defined separate models to predict SCNA scores and CYT from tumor stages, with comparison to Stage I as a baseline (Table 1, see Methods). In the model predicting SCNA scores, we observed significant positive coefficients (p= 1.39e-09, p= 3.77e-10, p= 2.01e-11) for each tumor stage (Table 1). For CYT, we observed almost significant negative coefficients for Stage II (p = 0.075) and a significant negative coefficient for Stage IV (p = 9.74e-5) (Table 1).

**Table 1.**
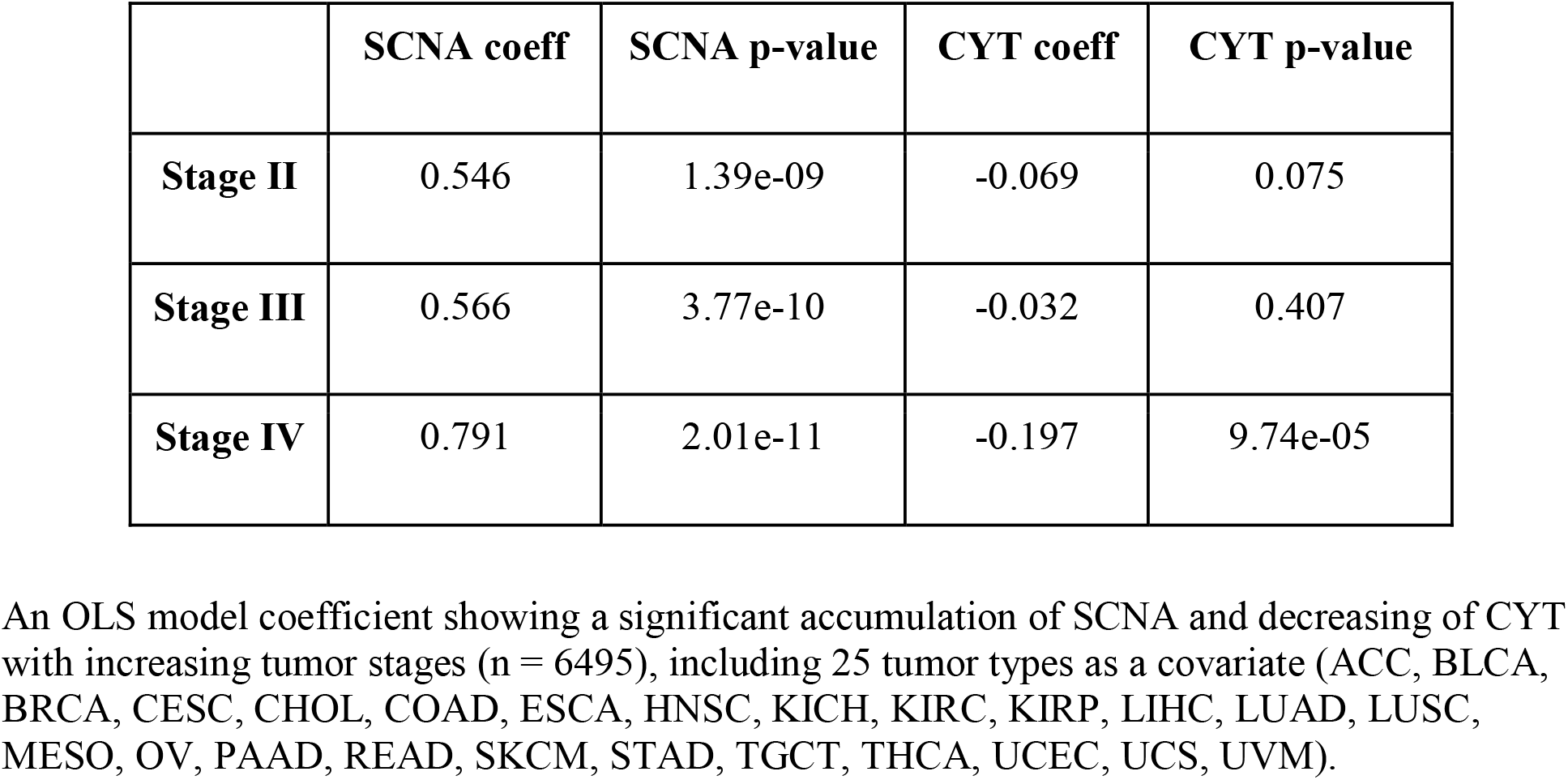
Significant accumulation of SCNA correlates with decreasing of CYT with tumor stage progression.

|  | SCNA coeff | SCNA p-value | CYT coeff | CYT p-value |
| --- | --- | --- | --- | --- |
| **Stage II** | 0.546 | 1.39e-09 | -0.069 | 0.075 |
| **Stage III** | 0.566 | 3.77e-10 | -0.032 | 0.407 |
| **Stage IV** | 0.791 | 2.01e-11 | -0.197 | 9.74e-05 |

An OLS model coefficient showing a significant accumulation of SCNA and decreasing of CYT with increasing tumor stages ( $n = 6495$ ), including 25 tumor types as a covariate (ACC, BLCA, BRCA, CESC, CHOL, COAD, ESCA, HNSC, KICH, KIRC, KIRP, LIHC, LUAD, LUSC, MESO, OV, PAAD, READ, SKCM, STAD, TGCT, THCA, UCEC, UCS, UVM).

**Figure 2.**
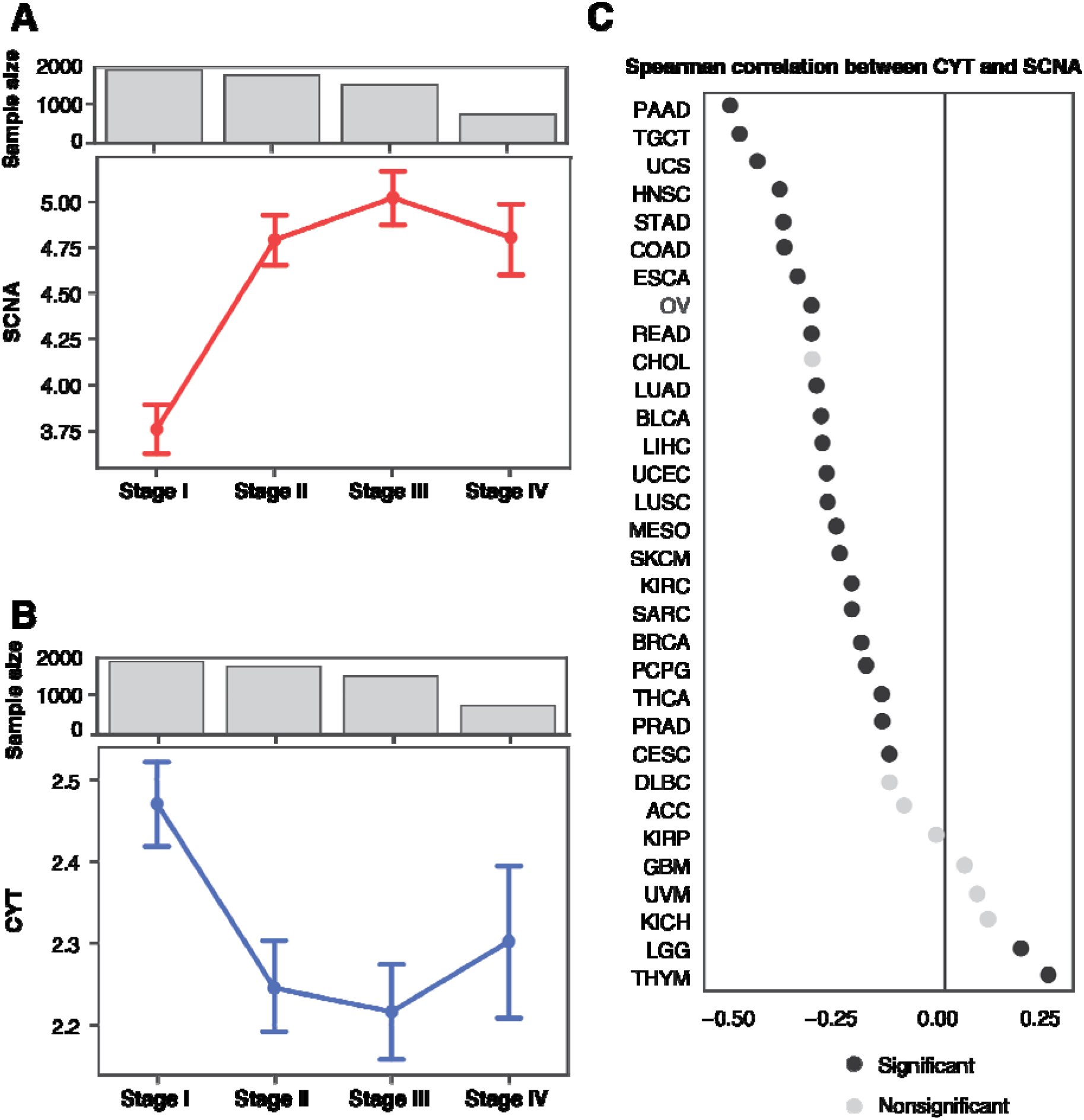
SCNA accumulates as tumor stage progresses and negatively correlates with immune cytolytic activity. (**A**) Mean and 95% confidence interval is shown for SCNA scores among samples at each tumor stage across 6298 TCGA samples with stage annotation and SCNA score available. (**B**) Mean and 95% confidence interval for CYT scores among samples at each tumor stage across 6458 TCGA samples with stage annotation and CYT score available. (**C**) Spearman correlation coefficients linking SCNA and CYT scores across 32 tumor types. Black circles denote significant correlation (FDR < 0.05) after Benjamini–Hochberg multiple testing correction.

We found significant negative correlation between SCNA scores and CYT levels in twenty-three out of thirty-two tumor types (Spearman correlation test; Fig. 2C). Surprisingly, low grade glioma (LGG) and thymoma (THYM) showed a significant positive correlation between SCNA score and CYT (Fig. 2C). Together, these observations suggest that as tumors progress, they accumulate SCNAs and evade immunity.

A previous report suggested that among SCNA categories, whole chromosome and arm level event burden are predictive of immune evasion whereas focal event burden are associated with cell cycle (Davoli et al., 2017). We found that focal SCNAs also inversely correlates with CYT levels (Fig. S3B,C,D), albeit more weakly than chromosome or arm level SCNAs (Fig. S4A) in an OLS model using all 3 categories of SCNA to predict CYT score and including tumor types as a covariate. This supported the use of the combined SCNA score to analyze the impact of chromosomal abnormalities during tumor progression on immune dysregulation (i.e., decrease cytolytic activity) leading to progressive immune incompetence and immune evasion. In considering the effects of SCNA levels on UPR maintenance of proteostasis, we also did not expect the three SCNA categories to substantially differ from one another. Indeed, they all showed similar Spearman correlation to parental UPR gene expression across all 32 tumor types (Fig. S4B). In light of this, all subsequent analyses were performed using the single SCNA score as a simplified measure of aneuploidy burden.

### UPR gene expression correlates with SCNAs

The UPR is an adaptive survival mechanism used by mammalian cells in response to environmental perturbations, cell-autonomous and cell-nonautonomous signaling to alleviate the burden of excess client proteins in the ER (Walter and Ron, 2011). To investigate the relation between SCNA and the UPR, we first examined the expression of a few representative genes from each major UPR pathway. We compared gene expression levels for the master regulator of the UPR, Heat Shock Protein Family A Member 5 (*HSPA5*) in tumors and matched normal tissues. Out of the twenty-three tumor types with available matched normal samples in TCGA, all except three (THCA, KICH, and KIRP) showed greater *HSPA5* expression in tumors, and thirteen of these showed statistical significance (FDR < 0.05) (Fig. 3A). Notably, small sample sizes for matched normal tissues limited the statistical power in a few cancer types: skin cutaneous melanoma (SKCM, n = 1), thymoma (THYM, n = 2), and pheochromocytoma or paraganglioma (PCPG, n = 3). We next evaluated the Spearman correlation between SCNA score and parent genes for the three branches of the UPR (IRE1α, PERK and ATF6) across all thirty-two tumor types in TCGA (Fig. 3B). Three genes from the PERK pathway (*EIF2S1*, *EIF2AK3*, and *DDIT3*) showed a positive correlation with SCNA score across almost every tumor type. *ATF6* also showed a mild positive correlation with SCNA scores across the majority of tumor types (Fig. 3B). In contrast *ERN1* (the gene coding for IRE1α) showed no consistent correlation, and *XBP1* had a mild negative correlation with SCNA score (Fig. 3B). This analysis of transcriptional regulation of sensor genes suggested, therefore, that SCNA levels correlate with activation of UPR branch pathways, mainly the PERK pathway. A positive correlation with *ATF6* is not entirely surprising given its role in targeting stress response genes to cope with a greater client protein burden resulting from SCNAs and facilitating tolerance to chronic stress (Wu et al., 2007). On the other hand, the lack of a positive correlation with *ERN1* motivated further analysis given that this pathway has been implicated in tumor survival (Logue et al., 2018; Xie et al., 2018), macrophage polarization (Batista et al., 2020), and T-cell dysregulation (Song et al., 2018).

**Figure 3.**
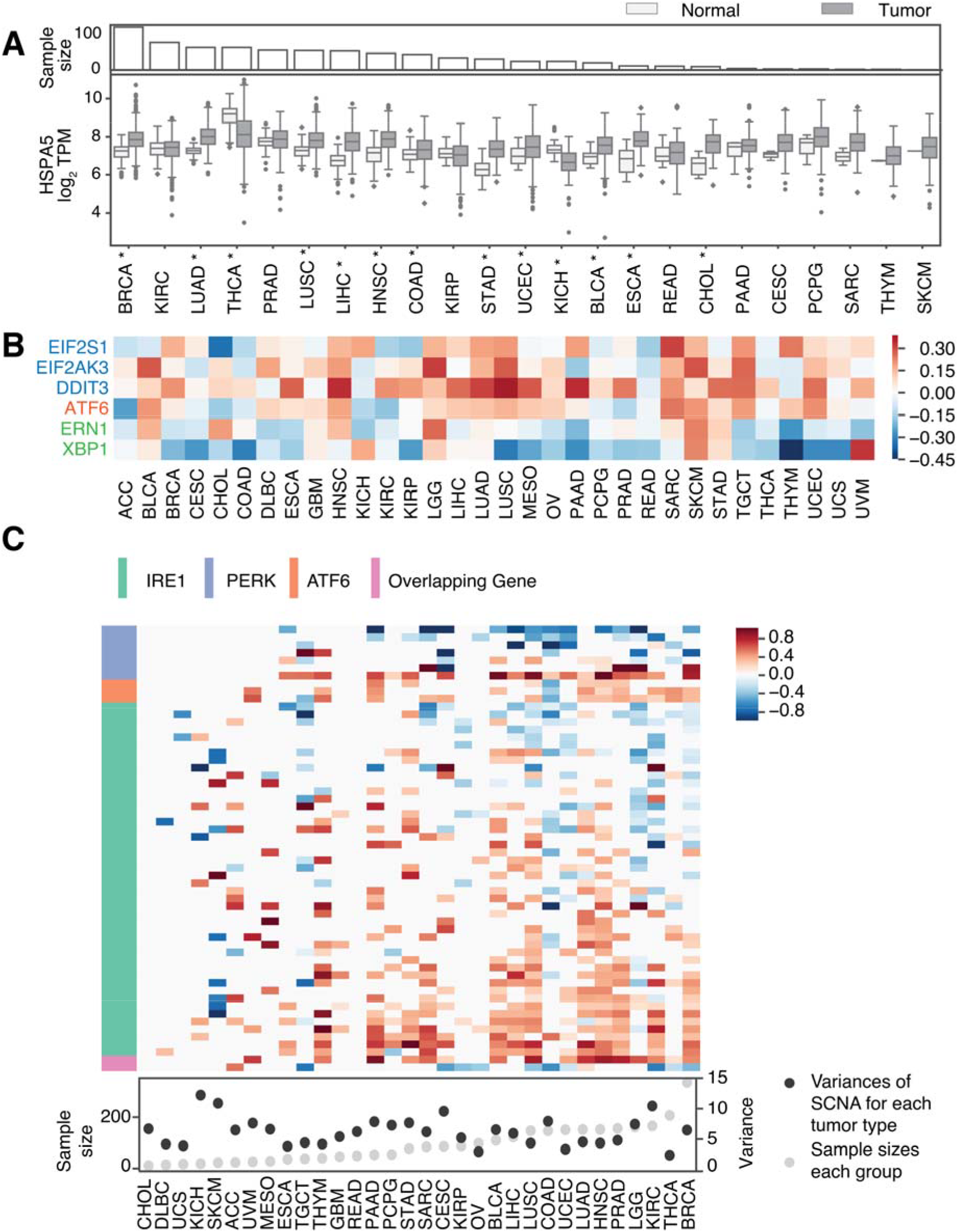
The unfolded protein response is influenced by SCNA levels. (**A**) A boxplot showing log_2_ TPM values of HSPA5 gene expression compared between tumors and matched normal samples across 23 tumor types from TCGA with normal tissue data available. The barplot on the top shows the number of normal tissue samples for each corresponding tumor type. Asterisks indicate significant differences by student’s t-test after Benjamini–Hochberg multiple testing correction (FDR < 0.05). (**B**) A heat map showing the Spearman correlation coefficient between the log_2_ TPM expression of three UPR branch pathway parental genes (rows) and SCNA scores across 32 tumor types (columns). Red cells represent positive correlation coefficients (r > 0) and blue cells represent negative correlation coefficients (r < 0). Gene names colored in blue belong to the PERK pathway, orange to the ATF6 pathway and green to the IRE1 pathway. (**C**) A heatmap showing the differential expressions of all genes from the three branches of UPR between low and high SCNA groups across tumor types. Low and high SCNA status was defined as the bottom and top 30% of samples within each tumor type. Rows depict genes from the UPR branch pathways from REACTOME and columns depict the 32 tumor types. Cells are colored in red or blue if the gene showed significant differential expression between low and high SCNA groups in that tumor type (Wilcoxon rank-sum test after Benjamini-Hochberg multiple hypothesis correction, FDR < 0.05). Color intensity corresponds to the log_2_ fold change in expression. The left side bar indicates pathway membership of the genes. The bottom panel shows the variances of SCNA scores for each tumor type and the number of normal tissue samples available for differential expression analysis.

Some UPR activity, and IRE1α activity in particular, is regulated by post-translational modifications which may not be reflected in expression levels of UPR branch pathway genes. Based on this reasoning, we performed an analysis of genes downstream of each of the three main branches of the UPR, assuming that they would collectively be more indicative of an association with SCNA levels than the parent genes. We first collected gene sets for the IRE1α, PERK and ATF6 pathways from REACTOME, a curated database of biological pathways (Jassal et al., 2020) (Table S1). We then compared the expression of these downstream genes in each UPR branch in tumor samples with either high (> 70% quantile) or low (< 30% quantile) SCNA score (Fig. 3C). Of note, inadequate coverage of samples and insufficient variation in SCNA levels posed limitations in this analysis (Fig S5). For example, CHOL, DLBC, UCS, KICH, ACC, and MESO all have particularly low numbers of samples in each group (n < 26, with mean sample size = 91.28), while GBM, OV, CESC, and THCA all have little variation in SCNA levels (variance < 5.41, with mean variance among all tumor types = 6.19). Despite this, we still found that over half of the thirty-two tumor types showed significant correlation between SCNA score and the expression of the majority of downstream genes in all three UPR branch pathways (Fig. 3C). Collectively, this broader analysis shows that SCNA is associated with perturbation of each of the three branches of the UPR.

### Changes in differential co-expression of UPR genes between SCNA^low^ and SCNA^high^ tumors

Next, we considered that UPR branch pathway activities themselves could be directly or indirectly affected by SCNAs. Because signaling requires the coordinated activity of multiple proteins, genes within pathways are often more highly co-expressed (Komili and Silver, 2008; Wolfe et al., 2005). Therefore, to assess the impact of SCNA levels on UPR signaling, we evaluated the differential co-expression of all UPR genes in low and high SCNA groups across tumor types. We used the same threshold to divide samples into SCNA^low^ and SCNA^high^ groups for each tumor type and assessed differences in the pairwise correlation coefficients for all UPR genes between these two groups. We found that almost universally the co-expression patterns of UPR genes were visibly different between SCNA^low^ and SCNA^high^ groups (Fig. S6), with most tumor types showing less co-expression in the SCNA^high^ compared to the SCNA^low^ group (Fig. 4A; Fig. S6) consistent with general perturbation of the transcriptome by SCNAs. In general, the SCNA^high^ condition showed loss of coordination of UPR genes relative to the SCNA^low^ condition (Fig. S6). The strongest effects were observed in PAAD, GBM, KICH, CHOL, UVM and ESCA.

**Figure 4.**
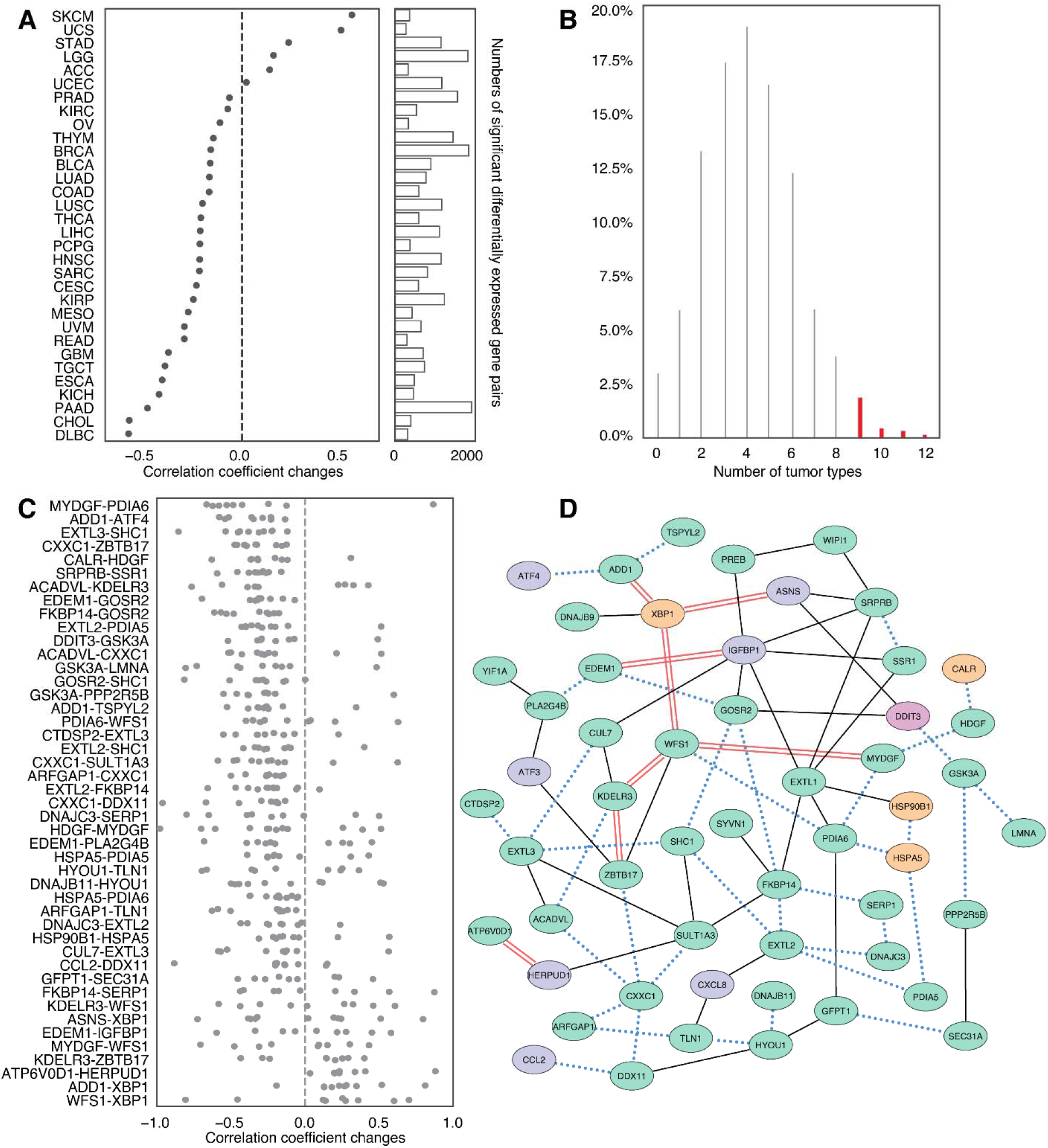
Co-expression analysis of UPR genes comparing low and high SCNA tumors. (**A**) A strip plot summarizing differences in co-expression (x-axis) of pairwise combinations of 58 UPR genes (n= 3364) between low and high SCNA groups across 32 tumor types (y-axis) as quantified by the change in Spearman correlation coefficient. The side bar indicates the number of gene pairs with significant co-expression change relative to a null distribution obtained from 1000 permutations of SCNA status (see Methods for detail). (**B**) A histogram showing the percentage of gene pairs (y-axis; n = 3364) that have significant co-expression change according to the number of types (x-axis) in which each gene pair was significant. Colored bars indicate the 2.68% of gene pairs (n = 45) that were significant in at least 9 tumor types, and that were selected for more in depth analysis. (**C**) Change in Spearman correlation coefficient between SCNA high and low conditions for 45 gene pairs with significant co-expression changes across more than 9 tumor types (n = 45). Each point indicates the difference in correlation for one tumor type where the gene pair was significant. (**D**) Network plot showing top UPR gene pairs with reduced, preserved or augmented co-expression. Each node represents a UPR gene and each edge represents a co-expression relationship between a gene pair. Red double line edges depict increased co-expression in SCNA^high^ tumors compared to SCNA^low^. Solid lines depict preserved co-expression between gene pairs and blue dotted lines depict reduced co-expression between gene pairs. Node colors represent the UPR branch pathway membership of genes, with green representing the IRE1 pathway, blue representing the PERK pathway, orange representing the ATF6 pathway and purple representing membership in more than one branch pathway.

We speculated that in the setting of SCNA^high^ the oncogenic effects of UPR are preserved or amplified while tumor suppressive aspects are reduced. We therefore evaluated whether loss of coordination of gene expression under SCNA^high^ conditions appeared random by comparing to permuted data. Interestingly, we observed that fourty-eight gene pairs showed a significant propensity to co-expression change (FDR <0.05) in at least nine tumor types (Fig. 4B). In these gene pairs, the co-expression changes were predominately negative (n = 37), suggesting a pattern of loss of coordination (Fig. 4C). Some genes were included in multiple perturbed pairs. Among highly perturbed gene pairs across multiple tumor types we noted *CXXC1, HSPA5, GSK3A, SERP1, PDIA6, FKBP14,* and *SCH1*, which showed multiple co-expression changes. Most of these genes (*HSPA5*, *CXXC1*, *SERP1*, *SCH1, PDIA6*) encode proteins that confer resistance to various forms of stress. GSK3A additionally functions as an oncogene by stabilizing β-catenin and promoting self-renewal. These genes have been associated with unfavorable prognosis in a cancer type related manner (Fig. S7) (Uhlen et al., 2017). On the other hand, co-expression of some gene pairs was preserved across all tumor types despite increased SCNA. We identified 34 gene pairs involving 35 genes that showed significant correlation in all tumor types (FDR < 0.05; Table S2). Gene ontology analysis of genes with reduced, augmented or preserved co-expression suggested that genes with preserved or augmented co-expression, but not those with perturbed co-expression, were associated with negative regulation of apoptosis (Fig. S8, Table S3, GO:1902236, GO:2001243, GO:2001234, GO:0043066).

We summarized perturbed, augmented and preserved gene co-expression relationships with a network (Fig 4D). This highlights more preserved relationships among genes involved in core activities of the ER such as cellular metabolism and co-translational translocation to the ER (IGFBP1-SRPRB, IGFBP1-SRR1) and pairs with at least one member involved in protein trafficking (PREB-WIPI1, KDELR3-CUL7, GOSR2-IGFBP1, SYVN1-FKBP14, YIF1A-PLA2G4B). We note that the relationship between ATF4 and DDIT3, while not consistently perturbed - ATF4 and DDIT3 co-expression was only significantly perturbed in 5 tumor types (Fig S9) - was also not preserved. Interestingly *DDIT3* co-expression with *GOSR2* (protein transport) and *ASNS* (asparagine synthetase) remained coordinated, suggesting that some less known aspects of DDIT3 activity may benefit tumor cells. DDIT3 is frequently thought of as the major executioner of apoptosis downstream of irrecoverable UPR stress, but it may be required for other functions, for example the induction of the proinflammatory/tumorigenic cytokine IL-23 (Goodall et al., 2010). While a deeper analysis of the coordinated activities of UPR proteins is merited, overall, this pattern is consistent with promoting positive aspects of UPR signaling. These may include regulation of metabolism, transport and bioenergetics, favoring cell survival while diminishing effects disadvantageous to the cell such apoptosis. Preservation of the UPR in SCNA^high^ tumors argues therefore for an active stress response to proteostasis. Since UPR signaling is known to affect immune cells, we next interrogated the UPR as the link between SCNA and reduced CYT.

### UPR activity links SCNA and CYT

Given that overall the UPR is activated in tumors relative to normal tissues (Fig 3C), but increasing SCNA levels make expression-based assessment of pathway activity from individual genes ambiguous, we developed a strategy to measure pathway activation from the combined effects of multiple genes. To establish a gene expression-based method for assessing UPR branch pathway activity in tumors, we adapted the pathway measurement method of Schubert et al (Schubert et al., 2018) applying a regression model to assign coefficients for genes within pathways and then deriving aggregate pathway activation scores by matrix multiplication. Whereas previous authors used a linear regression model to extract gene coefficients, we used a Lasso regression model to remove redundant genes from each pathway, to avoid overfitting and capture dominant differences (Fig. S10, Table S4). We applied this method using gene sets from REACTOME (58 genes) (Jassal et al., 2020) as previously described, further distinguishing IRE1α into its known functions, XBP1 splicing and RIDD, as these are non-overlapping activities. Our final scores represent differential activity in each UPR branch based on contrasting expression of genes in tumors and matched normal tissues (n = 23). Due to the limitation imposed by lack of matched normal tissues, we were only able to acquire pathway scores for twenty-three tumor types (see Methods).

Among UPR branch pathways, we found that the PERK pathway had a strong inverse correlation with CYT (Fig. 5A, n = 19 tumor types with non-zero pathway score). We then interrogated the IRE1α pathway by looking at XBP1, the canonical target of IRE1α endonuclease activity. The pathway score for spliced XBP1 (XBP1s) trended toward a mild negative correlation with CYT score and a positive correlation with SCNA (Fig. 5B, n = 18 tumor types with non-zero pathway score).

**Figure 5.**
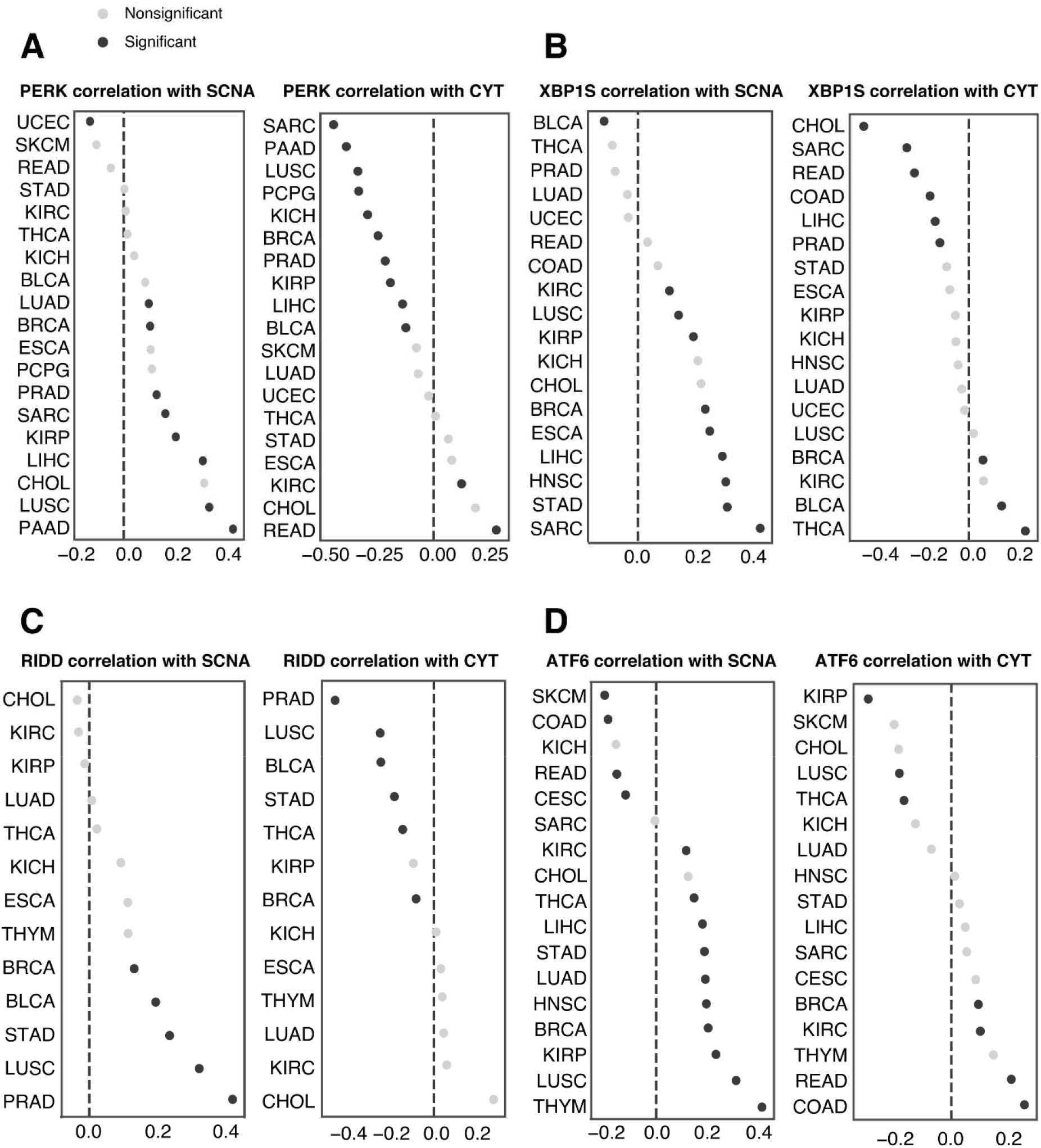
RIDD and PERK pathway activity scores show an inverse correlation with CYT score. (**A**) Spearman correlation coefficients (x axis) linking PERK pathway score with SCNA score (left), and CYT score (right) across 19 tumor types for which PERK pathway scores could be calculated. (**B**) Spearman correlation coefficients (x axis) linking XBP1S pathway score with SCNA scores (left), and CYT score (right) across 18 tumor types for which XBP1S pathway scores could be calculated. (**C**) Spearman correlation coefficients (x axis) linking RIDD activity score with SCNA score(left), and CYT score (right) across 13 tumor types for which RIDD activity scores could be calculated. (**D**) Spearman correlation coefficients (x axis) linking ATF6 pathway score with SCNA score (left), and CYT score (right) across 17 tumor types for which ATF6 pathway scores could be calculated.

The IRE1α pathway has a second downstream activity besides XBP1 splicing: regulated IRE1α -dependent decay of mRNA or RIDD (Hollien et al., 2009). Because of this functional duality, we decided to fully explore the signal from IRE1α by extracting RIDD’s thirty-three target genes (Maurel et al., 2014). The RIDD pathway score was both significantly positively correlated with SCNA and negatively correlated with CYT in five tumor types (BRCA, BLCA, STAD, LUSC, PRAD) (Fig. 5C). We observed largely positive correlation between ATF6 and SCNA level but little correlation with CYT score (Fig. 5D). Collectively, our analysis suggests that both IRE1α (through its RIDD activity) and PERK are associated with mechanisms of immune evasion in the tumor microenvironment.

We next evaluated the three UPR pathways relative to the effect of SCNA on the CYT score controlling for tumor type and purity in a single model. Tumor purity was included as a possible confounding factor since SCNA scores could be underestimated for lower purity tumors, and higher levels of immune infiltrate could inflate CYT scores. We obtained IHC-based estimates of tumor purity for TCGA from (Aran et al., 2015). We then applied an OLS linear model to evaluate the relative contributions of SCNA together with all UPR branches and IHC score in predicting CYT, including tumor type as a covariate, and limiting analysis to samples from the sixteen tumor types for which IHC scores were available. We found that SCNA had a highly significant negative coefficient (coefficient = −0.302, p<1.19e-118) in predicting CYT (Table 2). Similar to SCNA, both RIDD and PERK had negative coefficients (RIDD coefficient = −0.035, p > 0.471; PERK coefficient = −0.274 p < 3.61e-10), though only PERK was significant, suggesting that these UPR branches are associated with reduced immune activity. In contrast, ATF6 had a positive effect on CYT levels (coefficient = 0.238, p < 2.28e-5) and XBP1s was not associated with CYT scores (coefficient = 0.064, p > 0.147). In a model without IHC, RIDD reached statistical significance (coefficient = −0.142, p < 0.010, Table S5), pointing to infiltrating immune cells as the source of CYT suppressive RIDD signaling. XBP1s activity remained unassociated (coefficient = 0.092, p < 0.062).

**Table 2.** Coefficient of an OLS regression model using UPR pathway scores and SCNA scores.

|  | **Coeff** | **p-value** | **95% CI** |
| --- | --- | --- | --- |
| **SCNA** | -0.302 | 1.19e-118 | -0.328, -0.277 |
| **XBP1S** | 0.064 | 0.149 | -0.023, 0.150 |
| **PERK** | -0.274 | 3.61e-10 | -0.359, -0.188 |
| **ATF6** | 0.238 | 2.28e-05 | 0.128, 0.348 |
| **RIDD** | -0.035 | 0.471 | -0.131, 0.060 |
| **IHC** | -0.094 | 8.23e-17 | -0.116, -0.072 |

Coefficient of an OLS regression model using 4 UPR pathway scores and SCNA scores, including 16 tumor types (ACC, BLCA, BRCA, CESC, COAD, GBM, HNSC, KICH, KIRC, KIRP, LGG, LIHC, LUAD, LUSC, OV, PRAD, READ, SKCM, THCA, UCEC, UCS) with IHC data available as covariate to predict CYT, n = 7802.

Lack of correlation with XBP1 in tumor cells is not surprising given the demonstration that XBP1 in immune cells (dendritic cells and T cells) plays a tumor promoting role (Cubillos-Ruiz et al., 2017), hence highlighting the relevance of cell types and lineages in defining the role of UPR branches in the tumor microenvironment. Remarkably, RIDD genes expression was more suppressed in infiltrating macrophages than in tumor cells in single cell RNA expression data from (Tirosh et al., 2016) (Fig. S11A, B), in agreement with recent findings in murine macrophages (Batista et al., 2020). Collectively, the fact that RIDD and PERK have a similar relationship to CYT is not surprising since RIDD activity was shown to be PERK dependent (Moore and Hollien, 2015). To assess this dependent relationship, we evaluated the Spearman correlation between the PERK pathway and RIDD across the same sixteen tumor types. We found significant positive correlation in eight out of twelve tumors types (Fig. S12) where both RIDD and PERK pathway score were available, supporting functional interdependence. Thus, we conclude that among the UPR branch pathways, PERK and RIDD exert a negative effect on immune cells in the tumor microenvironment cooperatively.

### Experimental aneuploidy induces the UPR

A mechanistic link between aneuploidy and the UPR in cancer cells was sought using Reversine (Rv), a small molecule known to induce aneuploidy through inhibition of the mitotic spindle (Santaguida et al., 2017; Santaguida et al., 2015). To maximize the effect of Rv, we used two human cancer cell lines reported to be “quasi-diploid”: DLD1 (colon cancer; 2n= 46) and SKOV3 (ovarian cancer; 2n= 46) (Buick et al., 1985a; Knutsen et al., 2010). Digital karyotyping was performed as previously described (D’Antonio et al., 2017). In untreated DLD1 cells that present trisomy on 11p, Rv treatment promoted additional abnormalities (trisomy of 11q and of chromosome 20) (Fig. S13). We treated semi-confluent cells with varying concentrations of Rv for up to 72 hours and measured *XBP1* mRNA splicing by PCR as an indicator of an ER stress response (Fig. 6A). After treatment, both cell lines showed demonstrable ER stress with varying kinetics. A quantification of *XBP1* splicing reveals that maximal effects in DLD1 occurred at 12 hours while in SKOV3 at 72 hours (Fig. 6B). This shows that both cell lines respond to short-term Rv treatment activating the UPR, albeit with slightly different kinetics.

**Figure 6.**
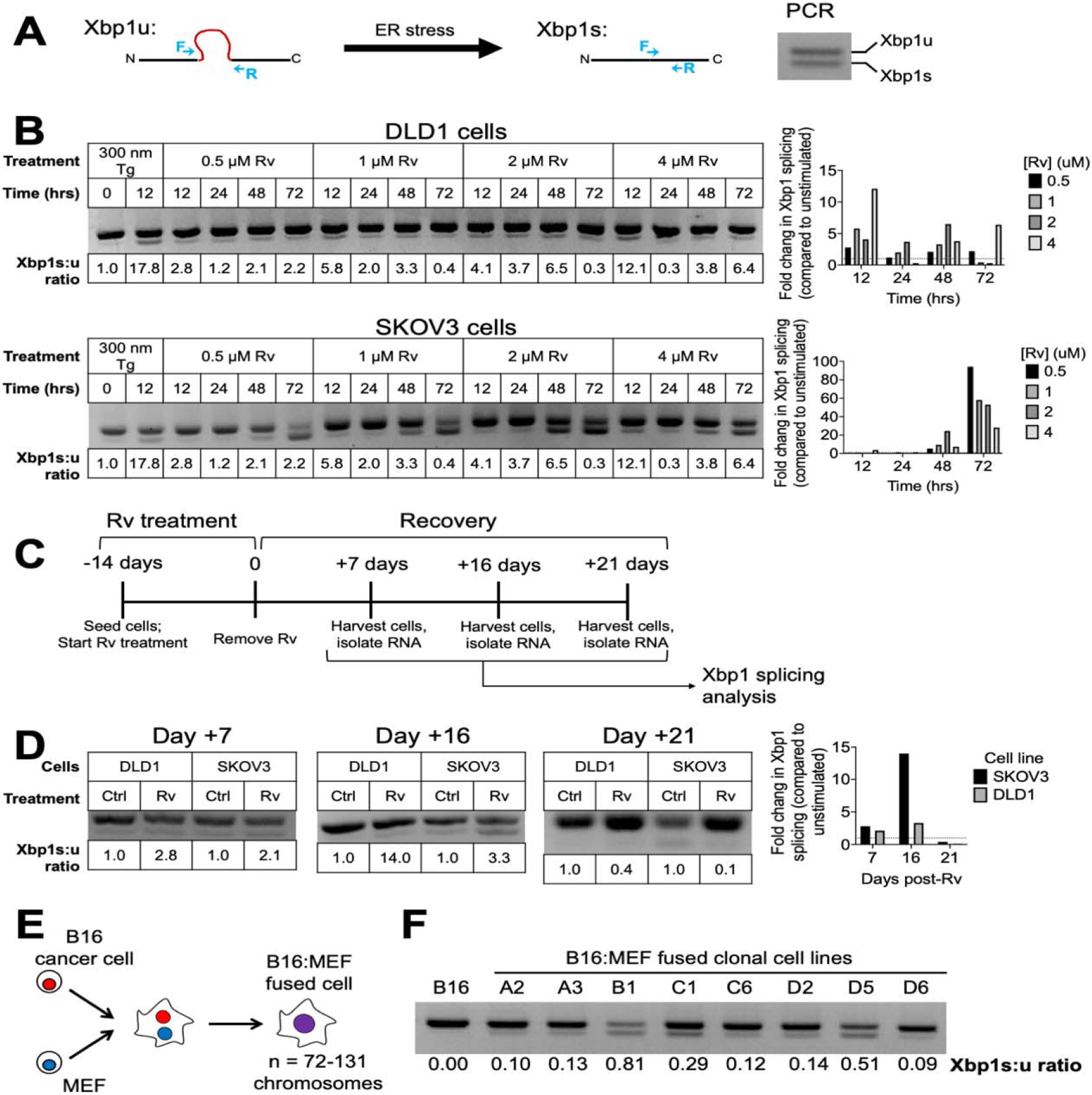
Experimental aneuploidy triggers XBP1 splicing in cancer cells. (**A**) Schematic representation depicting PCR-based analysis of Xbp1 splicing. During conditions of ER stress, a 26 base pair fragment is spliced from Xbp1 mRNA. To detect this, forward and reverse PCR primers (shown in blue) were designed to span the splice site. PCR amplification distinguishes between unspliced (Xbp1-u, upper band) and spliced (Xbp1-s, lower band) Xbp1 mRNA. To quantify ER stress, a ratio of spliced: unspliced Xbp1 was calculated. (**B**) Xbp1 splicing analysis and quantification of DLD1 and SKOV3 cells treated with varying concentrations of Rv for 12, 24, 48, and 72 hours. Thapsigargin (Tg) was used as a positive control. (**C**) Schematic representation for the work flow for the long-term Rv treatment experiment. (**D**) Xbp1 splicing analysis and quantification of DLD1 and SKOV3 cells treated with varying concentrations of Rv for 14 days and then recovered for 7, 16, and 21 days. (**E**) Schematic representation depicting cell-cell fusion between a B16 melanoma cell and a mouse embryonic fibroblast (MEF). (**F**) Xbp1 splicing analysis for B16 melanoma cells and eight B16:MEF fused clonal cell lines.

To determine if the effect of Rv on *XBP1* splicing was transient or sustained, we performed a second experiment with “long-term” Rv exposure (14 days) followed by a wash-out period (no Rv) for up to 3 weeks (Fig. 6C). We found that both DLD1 and SKOV3 cells had a sustained ER stress response for up to 16 days after Rv removal; by day 21 XBP1 splicing was no longer detected (Fig. 6D). Thus, prolonged treatment with Rv induces a UPR lasting several weeks after Rv removal linking aneuploidy and UPR both in acute and chronic conditions.

We sought independent validation by testing a model of non-pharmacologically induced aneuploidy. We used a panel of eight clonal cell lines derived through cell-cell fusion between B16 melanoma cells and mouse embryonic fibroblasts (MEF) (Searles et al., 2018) (Fig. 6E). The chromosome numbers in these fused cell lines range from 72-131 (Searles et al., 2018). We tested *Xbp1* mRNA splicing in each of the fused clones at baseline and compared it to the parental B16 cell line to see if fusion-driven aneuploidy induces the UPR. All (8/8) fused cell lines had higher amounts of *Xbp1* spliced isoform compared to unfused B16 cells (Fig. 6F). Thus, two independent models of experimental aneuploidy - Rv treatment and cell-cell fusion - both point to a mechanistic link between aneuploidy and UPR induction.

### Aneuploid cells polarize bone marrow-derived macrophages

Previously, we demonstrated that the conditioned media (CM) of ER stressed cancer cells polarizes macrophages and dendritic cells to a pro-inflammatory/immune-suppressive phenotype, impairing antigen-specific T cells (Mahadevan et al., 2012; Mahadevan et al., 2011). Subsequently, we demonstrated that these effects are operational *in vivo* and contribute to tumor development in an IRE1α-dependent manner (Batista et al., 2020). The present TCGA analysis showed an inverse correlation between single SCNA score and CYT across disease stages, suggesting that tumor cells with experimentally-induced aneuploidy could also dysregulate immune cells through a cell-nonautonomous mechanism. To this end, the CM of aneuploid cells collected at the time of maximal *XBP1* splicing was added to cultures of murine bone marrow-derived macrophages (BMDM) for 24 hours. We then isolated their RNA and analyzed the expression of a canonical pro-inflammatory cytokine (*Il6*) and the immune-suppressive enzyme Arginase 1 (*Arg1*) (Rodriguez et al., 2005). A schematic representation of the workflow for the experiment is shown in Fig. S14. Definitive *Xbp1* splicing was observed in BMDM treated with the CM of fused B16 cells but was only slightly by the Rv cell CM suggesting that the secretome of established aneuploid cells is more efficient at inducing a UPR in BMDM (Fig. 7A)

**Figure 7.**
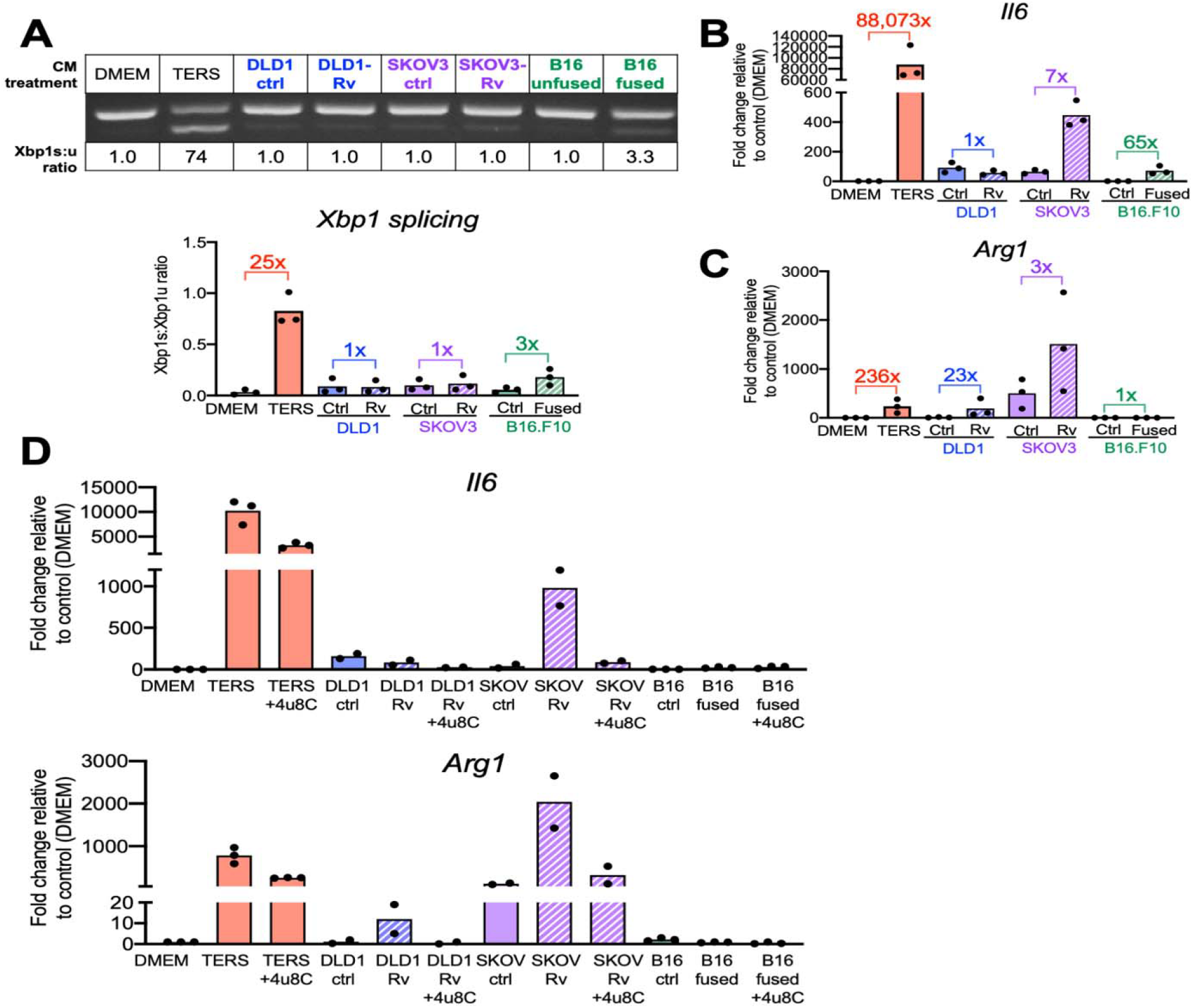
Cell nonautonomous effects on the aneuploid cancer cells on bone marrow-derived macrophages. (**A**) Xbp1 splicing analysis and quantification of bone marrow-derived macrophages (BMDM) cultured with the conditioned media of control- or Rv-treated DLD1 and SKOV3 cells, or unfused and fused B16 melanoma cells. (**B,C**) Quantification of *Il6* and *Arg1* mRNA in BMDM cultured with the conditioned media of control- or Rv-treated DLD1 and SKOV3 cells, and unfused and fused B16 melanoma cells via qPCR. (**D**) Quantification of Il6 and Arg1 mRNA in BMDM cultured with the conditioned media of control- or Rv-treated DLD1 and SKOV3 cells, and unfused and fused B16 melanoma cells in the presence or absence of the IRE1a inhibitor 4μ8C via qPCR.

Next, we looked at *Il6* gene expression, a pro-inflammatory/tumorigenic cytokine (Grivennikov et al., 2009). The CM of Rv-treated SKOV3 and fused B16 cells, but not DLD1 cells, yielded high *Il6* induction relative to controls (7-fold and 65-fold, respectively) (Fig. 7B). The CM from Rv-treated DLD1 and Rv-treated SKOV3 also yielded high *Arg1* expression levels compared to control cultures (23-fold and 3-fold, respectively) (Fig. 7C). Fused B16 cell CM was ineffective at inducing *Arg1*. Taken together these data demonstrate a functional link between aneuploidy, UPR, and the acquisition of a pro-inflammatory/immune-suppressive phenotype by macrophages.

Since the induction of *Il6* and *Arg1* in macrophages is under control of the IRE1α branch of the UPR (Batista et al., 2020) we sought to determine if their induction by the CM of aneuploid cells was also IRE1α dependent. We treated BMDM with the CM of Rv-treated or fused cells, with or without 4u8C, a small molecule inhibitor of IRE1α, and measured the transcriptional levels of *Il6* and *Arg1*. We found that in every instance where CM induced *Il6* and *Arg1* transcription, 4u8C markedly inhibited transcription (Fig. 7D), implying a direct involvement of IRE1α. Thus, experimentally-induced aneuploidy enabled us to establish a cause-effect relationship between cancer cell aneuploidy, the UPR and dysregulation of immune cells reminiscent of that observed in the tumor microenvironment.

## Discussion

Using a single SCNA score inclusive of whole-chromosome, arm and focal SCNA (aneuploidy burden) across 9,375 TCGA samples across 32 tumor types we provide evidence for an inverse correlation between SCNA and intra-tumor cytolytic activity (CYT), a proxy of local immune competence, in progressive stages of disease. In the same set of tumor samples we also found a correlation with the UPR, suggesting that the UPR is the likely link between aneuploidy and local immune dysregulation. In vitro models of aneuploidy induced pharmacologically via mis-segregation or cell-cell fusion provided a mechanistic validation.

Our hypothesis was that the UPR is the mechanism through which aneuploidy negatively affects local immunity (Zanetti, 2017). Our findings show that UPR gene expression correlates positively with aneuploidy, with genes of the PERK pathway showing strong positive correlation with SCNA across almost every tumor type. The first step in PERK’s homeostatic role in response to stress is the phosphorylation of eIF2α (eIF2α-P), which in turn inhibits global translation to attenuate the impact of client proteins inside the ER. Therefore, a positive correlation between SCNA and PERK, but not the IRE1α -XPB1 axis, is not surprising, even though in glioblastoma multiforme IRE1α has been shown to drive tumorigenicity (Lhomond et al., 2018). Since the UPR is an adaptive response, it follows that aneuploidy tolerance predisposes to an adaptive UPR, which then heightens cellular fitness and dysregulates local immune cells. Specifically, PERK engagement in tumor promotion can be a response to cell-autonomous (Bi et al., 2005; Hart et al., 2012) as well as cell-nonautonomous (Rodvold et al., 2017) stress signals. We previously showed that the latter enable cancer cell survival and drug resistance with contextual reduction of ATF4 and CHOP activation downstream of eIF2α (Rodvold et al., 2017) even though, paradoxically, ATF4 and CHOP can both mediate apoptosis under condition of acute stress (Hiramatsu et al., 2020; Oyadomari and Mori, 2004).

How cancer cells avoid apoptosis in favor of higher fitness is poorly understood. eIF2α-P is a convergence point of the UPR and the integrated stress response (Ron, 2002; Walter and Ron, 2011), and regulates the translation of molecules relevant to immune dysregulation and tumorigenicity. For instance, eIF2α-P post-translationally regulates PD-L1 expression in MYC transgenic/KRAS mutant murine tumor (Xu et al., 2019). Furthermore, eIF2α-P redirects the translation of 5’-untranslated regions (5’-UTRs) (Sendoel et al., 2017). In addition, to corroborate this interpretation, the data on co-expression presented here point to increased negative regulation of the apoptotic program together with an enhancement of metabolic/bioenergetic fitness of the cell. It is therefore temping to speculate that through the induction of the UPR, aneuploidy regulates the translational machinery of the cancer cell in a more complex way than just through a gene dose effect. For instance, studies in preneoplastic cells show that eIF2α-P and inactivation can direct the translational machinery towards eIF2A-dependent uORF translation and increased ribosome occupancy of 5’-UTRs augmenting protein synthesis (Sendoel et al., 2017). The extent to which this phenomenon is exploited by SCNA will need future exploration.

A weak negative correlation between SCNA and IRE1α does not preclude an involvement of IRE1α in response to aneuploidy. In fact, we found that the IRE1α-dependent RIDD activity correlates positively with SCNA and negatively with CYT in several tumor types. RIDD activity degrades target mRNAs selectively halting the production of proteins (Hollien and Weissman, 2006) and fulfills a function somehow complementary to that of eIF2α-P in that both regulate proteostasis to diminish the workload of client proteins in the endoplasmic reticulum during times of stress (Maurel et al., 2014). Therefore, it is not surprising that RIDD activity and PERK are linked functionally nor that RIDD has been found to be PERK-dependent - the depletion of PERK inhibits RIDD in a substrate-specific manner (Moore and Hollien, 2015). Also of interest is the emerging aspect of RIDD-degradation of miRNAs (miR17, miR34a, miR96, miR125b, and miR200) (Upton et al., 2012; Wang et al., 2018). Among them, miR34a has been shown to exert negative regulation on PD-L1 expression (Wang et al., 2015), induce cellular senescence via modulation of telomerase activity (Xu et al., 2015), and inhibit adrenergic transdifferentiation of tumor associated sensory nerves in oral cancers in a p53-dependent manner (Amit et al., 2020). Thus, the effect of aneuploidy on RIDD activity introduces a new dimension in our understanding of intratumor immune dysregulation and tumorigenicity. In light of the above considerations we propose that aneuploidy coordinates two apparently distinct UPR pathways: PERK (through eIF2α) and IRE1α (through RIDD), which work interdependently to oppose local immune surveillance trough diminished cytolytic activity of intratumor T cells and dysregulation of macrophages and dendritic cells.

Cell-nonautonomous signaling through the UPR or the integrated stress response has been well documented in *C. elegans* and shown to increase longevity but also establish a neuroimmune axis of communication (Frakes et al., 2020; O’Brien et al., 2018; Taylor and Dillin, 2013; van Oosten-Hawle et al., 2013). A similar type of UPR-based transcellular communication has been documented in mammalian cells and specifically between cancer cells and bone marrow-derived myeloid cells (macrophages and dendritic cells) (Cubillos-Ruiz et al., 2015; Mahadevan et al., 2012; Mahadevan et al., 2011; Rodvold et al., 2017). The implication of transcellular communication is relevant not only to the biology of cancer cells, but also to the immunobiology of intra-tumoral macrophages and dendritic cells. While the nature of the transmitting factor(s) has remained elusive in all reports, the phenomenon is clearly relevant *in vivo*. In both *C. elegans* and mammalian cells the phenomenon is dependent on the IRE1α - XBP1 axis in receiver cells (Batista et al., 2020; Cubillos-Ruiz et al., 2015; Frakes et al., 2020; Taylor and Dillin, 2013). In macrophages and dendritic cells, an unintended consequence of transcellular communication of UPR signaling is the acquisition of a pro-inflammatory/immune suppressive phenotype, which is also found in tumor-bearing mice (51) and in cancer patients (Chittezhath et al., 2014; Sousa et al., 2015). Here we used two *in vitro* models to show that aneuploidy as a sole trigger of the UPR (Fig. 6) is sufficient to modulate the phenotype of machophages in a cell-nonautonomous way. The experiments (Fig. 7) unequivocally show the specific transcriptional activation of *Il6,* a prototype pro-inflammatory/tumorigenic cytokine gene, and *Agr1*, a gene coding for a key T cell suppressive enzyme. Furthermore, we found that both *Il6* and *Arg1* transcription was markedly diminished by a small molecule inhitior of IRE1α RNAse activity (Fig. 7) consistent with the fact that this type of cell-nonautonomous regulation of myeloid immune cells is IRE1α-XBP1 dependent (Batista et al., 2020; Cubillos-Ruiz et al., 2015). Thus, the secretome of aneuploidy cells can remodel the phenotype of macrophages and dendritic cells as described previously *in vitro* and *in vivo* (Cubillos-Ruiz et al., 2015; Mahadevan et al., 2012; Mahadevan et al., 2011; Rodvold et al., 2017), *51*). Collectively, the data add a new layer of complexity to our understanding of the origin of immune dysregulation in the tumor microenvironment. If in fact signals emanating from aneuploid cells impart a pro-tumorigenic phenotype to macrophages and dendritic cells, focus should be placed on blocking community effects rather than cognate cell-cell interactions. For instance, establishing the role of IRE1α in macrophages and dendritic cells isolated from human cancers should be prioritized as this could lead to a new therapuetic angle to subvert local immune dysregulation. At a more general level it will be important to develop models to study the physico-spatial characteristics of transcellular communication between cancer cells and myeloid cells in the tumor microenvironment.

An unanswered question raised by the present study is *when* aneuploidy exerts its effects on the UPR relative to tumor history. It is known that aneuploidy increases during tumor evolution (Ben-David and Amon, 2020; Newburger et al., 2013) and correlates with poor prognosis (Owainati et al., 1987; Stopsack et al., 2019). Here we show that aneuploidy accumulates over the life of the tumor from stage I through stage IV (Fig. 2A). As shown, SCNA^high^ tumors differ drastically in gene co-expression patterns relative to SCNA^low^ tumors, suggesting that SCNA drives loss of connectivity among genes (Fig. 4C). Compared to other genomic alterations timed to early cancer evolution such as non-synonymous driver mutations (Vogelstein et al., 2013) and clustered mutational processes (chromothripsis) (Consortium, 2020), the impact of aneuploidy on the UPR may be stochastic and nonperpetual since advanced SCNA is associated with a marked loss of connectivity among UPR genes in the SCNA^high^ group across tumor types. Paradoxically, ovarian cancer, a tumor with the highest aneuploidy burden shows only a weak correlation with the UPR reflecting perhaps the fact that most women are diagnosed at stage III to IV, when the tumor has metastasized to the peritoneum and distant organs. Collectively, our data suggest that the effects of aneuploidy on the UPR during cancer evolution may be a progressive and possibly cumulative event until loss of gene connectivity takes place. However, we believe that once set in motion the unintended consequences on neighboring immune cells, i.e., loss of cytolytic activity and dysregulation of macrophages and dendritic cells, will persist, hampering natural immune surveillance and response to immunotherapy.

## Conclusions

Cancer comprises in excess of 100 different disease entities with diverse risk factors and epidemiology (Alexandrov et al., 2013; Tomasetti et al., 2017). Aneuploidy is an early event that determines genomic instability (Duesberg et al., 1998), increases substantially with stage of progression, and is associated with poor prognosis (Hieronymus et al., 2018). Its relationship with immunity has recently emerged with some reports suggesting that aneuploid cells are targeted by T and NK cells (Santaguida et al., 2017; Senovilla et al., 2012), and others showing that aneuploidy is inversely correlated with immune evasion (Davoli et al., 2017; Taylor et al., 2018). The hypothesis tested here demonstrates that a single SCNA score encompassing whole-chromosome, arm, and focal aneuploidy is sufficient to establish a positive correlation with the UPR and an inverse correlation with intratumor T cell immunity. Relevantly, we demonstrate that aneuploidy triggers the UPR which we identify as the mechanism through aneuploid cells remodel the immune cell landscape of the tumor microenvironment. Since these UPR-based effects can be propagated transcellularly with no need for cognate cell-cell interactions it appears as though immune dysregulation in the tumor microenvironment is the consequence of a community effect in which the UPR serves as the key mechanism enabling aneuploid cells to alert and modify the immune microenvironment. Current immune checkpoint blockade therapies are still ineffective in the majority of cancer patients and methods to predict likely responders are critically needed. Among current predictors, the tumor mutational burden is an imperfect indicator of responsiveness and DNA hypomethylation that correlates inversely with immune evasion also shows a direct correlation with SCNA (Jung et al., 2019). Here we propose that aneuploidy plays both a role in driving tumor adaptive evolution by providing fitness advantage to cancer cells (Pavelka et al., 2010) and in initiating/amplifying immune cell dysregulation and immune evasion. Therefore, standardized methods (Douville et al., 2018) to assess aneuploidy burden on an individual basis could help better stratify patients likely to respond to immune checkpoint blockade therapies. Compared to the tumor mutational burden, the aneuploid burden provides in addition insights into the degree of erosion of intra-tumor immune surveillance.

## Author contributions

Original Concept, M.Z.; Project Supervision, H.C. and M.Z.; Project Planning and Experimental Design, H.C., S.X., S.C.S. and M.Z.; Statistical Advising, T.C.W.; In vitro experiments, S.C.S., P.S.; Data Acquisition, Processing, and Analysis, S.X., S.C.S., K.J., A.C., and M.Z.; Data Analysis: S.X., S.C.S., K.J., A.C., and M.Z.; Writing Manuscript, H.C. and M.Z.

## Data and materials availability

Bioinformatic data have been deposited in https://github.com/cartercompbio/SCNA_score_analysis.

## Material and Methods

### Data

The TCGA files were downloaded from the gdc portal on 12/27/2017, using gdc-client v1.3.0. TCGA RNAseq alignment files were reprocessed using sailfish software version 0.7.4 and the GRCh38 reference genome with default parameters, and including all read sequence duplicates. Associated metadata were downloaded using TCGA REST API interface https://api.gdc.cancer.gov/files/. The MSI data were downloaded from (Kautto et al., 2017) supplementary data. We used a threshold of 0.4 as the cutoff for distinguishing MSI-H and MSS as suggested in this paper. Annotated somatic mutation calls from TCGA Pan-Cancer were downloaded from the GDC on 12/17/2016. TCGA Segmented SNP6 array data were downloaded from Broad Firehose (release stddata_2016_01_28, file extension: segmented_scna_hg19).

### Somatic copy-number alteration quantification

We considered three categories of SCNA as described previously (Davoli et al., 2017); whole chromosome, chromosome arm and focal copy number alterations. SCNAs were detected by comparing Affymetrix SNP data between tumor and paired normal samples. Based on the SNP intensity at the corresponding genomic position, we define a region as a contiguous set of SNPs with a shared log_2_ fold change in intensity. A region was designated an event if the log_2_ fold change exceeded certain thresholds. A log_2_ fold change greater than 0.1 or less than −0.1 was defined as a single event, and a log_2_ fold change greater than 1 or less than −1 as two events (Equation 1) (Beroukhim et al., 2010). Thus,

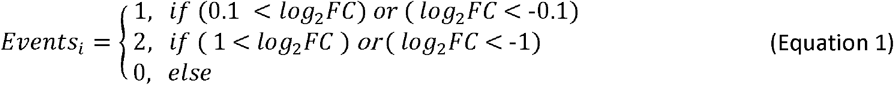

where *i* indexes regions of contiguous SNPs with the same intensity. Most regions are small, thus to score whole chromosome arms using Equation 1, we used the fractional length weighted sum of log_2_ fold changes across the regions within a chromosome arm *j* (Equation 2).

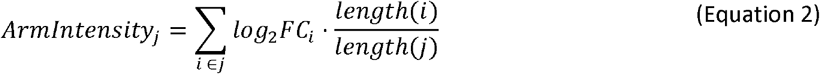

An event was designated whole chromosome if both chromosome arms met the Equation 1 criteria such that both arms were affected in the same direction, chromosome arm if one arm met the Equation 1 criteria or the arms were affected in different directions, and focal otherwise. Chromosomal and arm events were only counted once, in the largest category that applied. As focal events can happen subsequent to loss or gain of a chromosome or arm, we did not constrain counting of focal events based on the larger categories.

Events of each category were then summed for each sample. Whole chromosome and focal events were summed across 23 chromosomes, and chromosome arm level events were summed across 46 possible chromosome arms. As the resulting scores have very different ranges, (0-46 for chromosomal events, 0-92 for arm level events, and much larger values for focal events), we scaled each of these values before combining them into a single SCNA score (Equation 3) using sklearn.preprocessing.MinMaxScaler, with a feature range from 0 ~ 1.

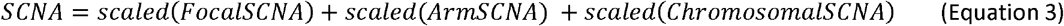

Since the FocalSCNA, ArmSCNA, and ChromosomalSCNA were all transformed to the same scale before aggregating, we interpret this SCNA score as a general reflection of genome abnormality, considering the 3 categories as contributing equally.

### Cytolytic activity

The Cytolytic activity (CYT) score was calculated as the geometric mean of log_2_ TPM expression values of granzyme A (*GZMA*) and perforin (*PRF1*) as described in (Rooney et al., 2015).

### SCNA correlation with non-silent mutation

We partitioned the TCGA samples into MSS (n = 8536) and MSI-H (n = 373) using the MSI data downloaded from (Kautto et al., 2017). We then removed silent mutations and computed the total number of mutations per sample. The relationship between the aggregated SCNA score and total number of non-silent somatic mutations was evaluated by Spearman correlation coefficient in MSS and MSI-H samples separately.

### *Tp53* mutations and P53 activity analysis

TCGA samples were partitioned into *TP53* wildtype and *TP53* mutated groups. Twenty five tumor types included at least one sample with *TP53* mutation (Figure S2A). The Wilcoxon rank-sum test was applied to test the aggregated SCNA score differences between *TP53* wildtype and *TP53* mutated groups within each tumor type (Figure S2A). P53 activity was calculated as the sum of z-scored log_2_ TPM expression values of 10 P53 downstream repressed genes, including *CCNB1, PLK1, EED, CDK1, EZH2, CCNB2, E2F3, MYBL2, FOXM1, E2F2* (Cancer Genome Atlas Research Network. Electronic address and Cancer Genome Atlas Research, 2017). Since P53 repression of these genes indicates P53 activity, the inverse of this value was used as the score representing P53 activity. This was done using sklearn.preprocessing.MinMaxScaler. The relationship between the P53 activity score and SCNAs score was assessed by Spearman correlation coefficient.

### OLS models fitting SCNA and CYT with tumor stages

An ordinary least square (OLS) linear model (Equation 4) was used to relate SCNA and CYT scores to tumor stage, including tumor type as a covariate to predict the independent variable using 6495 samples with stage information from 25 tumor types (ACC, BLCA, BRCA, CESC, CHOL, COAD, ESCA, HNSC, KICH, KIRC, KIRP, LIHC, LUAD, LUSC, MESO, OV, PAAD, READ, SKCM, STAD, TGCT, THCA, UCEC, UCS, UVM).

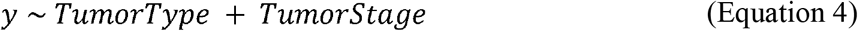

In Equation 4, *y* represents SCNA score or CYT score. Tumor stages and tumor types were encoded as categorical variables.

### Effects of SCNA score on UPR gene expression

TCGA samples were divided into 3 groups, SCNA^high^, SCNA^low^ and neither using the 30^th^ and 70^th^ percentiles of SCNA level within each tumor type. UPR gene expression levels were compared between SCNA^low^ and SCNA^high^ groups using the Wilcoxon rank sum test to determine whether there was a significant shift in expression between groups. Multiple hypothesis testing correction was performed using the Benjamini-Hochberg method with alpha = 0.05.

### Differential co-expression analysis of UPR pathway genes

Differential co-expression analysis was applied to test for pairwise co-expression changes between SCNA^high^ and SCNA^low^ samples (as defined above), using the method from (Tesson et al., 2010). First, the adjacency matrix for each phenotype was constructed by the following formula (Equation 5), where 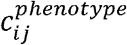 represents the Spearman correlation coefficient between gene *i* and *j* in a specified phenotype.

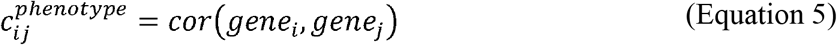

Then, the adjacency matrix difference is computed as follows (Equation 6), with the β parameter set to 4.

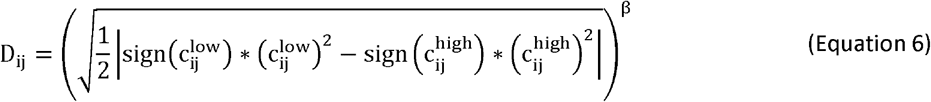

We then permuted SCNA group membership 1000 times within each tumor type to generate a null distribution for evaluating the significance of the pairwise correlation. This analysis included all 58 UPR genes, resulting in 3364 gene pairs. We identified gene pairs that showed less correlation than expected across more than 9 tumor types as recurrently perturbed, and pairs that showed more correlation than expected across all tumor types as preserved. Conserved gene pairs were further assessed by Spearman correlation pan-cancer, and only pairs that showed significant Spearman correlation (FDR < 0.05, multiple correction after Benjamini-Hochberg) were retained. The median co-expression change was calculated for each tumor type by summing the spearman correlation coefficient differences between SCNA^low^ and SCNA^high^ groups 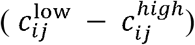 for each gene pair and taking the median (Fig. 4A). The number of gene pairs significant after permutation testing is shown in the side bar of Fig. 4A. The differences in correlation coefficient for selected gene pairs between SCNA^low^ and SCNA^high^ is shown in Fig. S6.

### GO Enrichment Analysis for selected gene pairs

We performed GO biological process analysis separately for 41 genes with recurrently perturbed co-expression patterns, 11 genes with augmented co-expression, and 35 genes with conserved co-expression patterns identified from differential co-expression analysis (above). GO Enrichment Analysis was performed using the online server http://geneontology.org/, using the “biological process complete” annotation data set with Homo sapiens reference list. The test result is calculated using Fisher’s Exact, with FDR cutoff < 0.05.

### UPR branch pathway score quantification

Gene sets representing PERK (Reactome id R-HSA-381042.1), XBP1s (Reactome id R-HSA-381038.2), and ATF6 (Reactome id R-HSA-381183.2) branch pathway downstream activity were extracted from the Reactome pathway database (Fabregat et al., 2018). The RIDD pathway downstream gene set was obtained from (Maurel et al., 2014). We implemented the pathway score quantification method from Schubert et al. 2018, however instead of applying non-regularized linear regression as in their work, we used Lasso regression to avoid overfitting and reduce redundant features. We built Lasso regression models using 10-fold cross validation to select the lambda parameter. In order to fit models that would represent the extent of induction of UPR branch pathways, we modeled the dependent variable using paired tissue-matched samples in TCGA such that *y* = 0 for tissue-matched normal samples and *y* = 1 for tumor samples. For each pathway, log_2_ TPM expression values of genes downstream of the branch pathway served as the independent variables. Models were fit in each tumor type separately. Matrix multiplication between the UPR branch gene expression matrix and the model coefficient matrix was applied to quantify pathway scores of individual pathways (XBP1s, PERK, ATF6 and RIDD) for each sample. Because the coefficient matrix represents the vector of corresponding genes in the plain of expression space, the pathway score is a meaningful representation of the distance from the origin (Schubert et al., 2018). Using this method, we obtained pathway scores for 7998 samples, across 23 tumor types that had normal tissue RNA-seq data available. Pathway scores were compared to SCNA and CYT scores by Spearman correlation.

### OLS model fitting UPR pathways, tumor types and SCNA to predict CYT

We fit an OLS model with XBP1s, PERK, ATF6, RIDD branch pathway scores, tumor type, tumor purity and SCNA scores as independent variables *x* to predict the dependent variable CYT score, *y* (Equation 7). Tumor purity was approximated by immunohistochemistry (IHC) measures obtained from (57). These data were available for 16 tumor types (n = 7802; 16 tumor types: BLCA, BRCA, CESC, COAD, HNSC, KICH, KIRC, KIRP, LIHC, LUAD, LUSC, PRAD, READ, SKCM, THCA, UCEC).

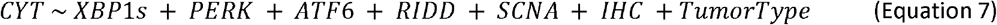

A second model was fit excluding IHC (n = 8488; 23 tumor types: BLCA, BRCA, CESC, CHOL, COAD, ESCA, HNSC, KICH, KIRC, KIRP, LIHC, LUAD, LUSC, PAAD, PCPG, PRAD, READ, SARC, SKCM, STAD, THCA, THYM, UCEC) using the formula

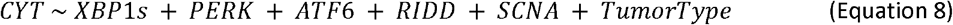

### Single Cell Analysis or RIDD pathway genes

We retrieved the single-cell data from (57) GSE72056. This dataset included measurements for 33 RIDD target genes. We performed hierarchical clustering of single cells using the 33 RIDD target genes with the Ward variance minimization algorithm using python package Scipy. We focused the analysis on tumor cells and macrophages as macrophages are the most abundant immune cell infiltrating tumors (Cassetta et al., 2019) and are involved in mediating cell non-autonomous effects that dysregulate the tumor microenvironment (Mahadevan et al., 2011) including RIDD activity (Batista et al., 2020). We excluded *ITGB2* and *TAPBP*, since *ITGB2* and *TAPBP* from further analysis, as they did not behave in the same way as other RIDD target genes (Batista et al., 2020). Mean expression of RIDD target genes was further compared using the Wilcoxon rank sum test.

### Software version, packages, and code availability

Computational analysis was performed using Python version 2.7.15. The OLS regression models used statsmodels.formula.api, version 0.9.0. Wilcoxon rank-sum tests, Spearman correlation analysis, hierarchical clustering and z-score calculations used scipy version 1.1.0. The LASSO regression model with cross-validation was applied using sklearn.linear_model.LassoCV, version 0.20.3. All rescaling was done using sklearn.preprocessing.MinMaxScaler, version 0.20.3. Plots were generated using matplotlib version 2.2.3, and seaborn version 0.9.0. Data representation used pandas version 0.24.2. Code and data to reproduce the analysis are available at <url to be determined>.

### Digital Karyotyping

Digital Karyotyping analysis was performed using Illumina Infinium Core-24 Beadarrays, which allow interrogation of >500,000 SNPs at single-nucleotide resolution. These arrays produce data from intensity signals corresponding to the presence of allele A and allele B at a given SNP. Using GenomeStudio (Illumina), we calculated the mean log R ratio, a measure of copy number as a ratio of observed to expected intensities; and the B-allele frequency, the proportion of allele calls at each genotype with respect to allele B (1.0 for B/B, 0.5 for A/B, and 0.0 for A/A). We created plots using these metrics to visually inspect each chromosome for abnormalities. For each kit we used 200 ng of DNA, which was processed according to manufacturer instructions. Following hybridization, BeadChips were scanned using the Illumina iScan System.

### Cell lines and culture conditions

The quasi-diploid cell lines DLD1 (colorectal adenocarcinoma) (Knutsen et al., 2010) and SKOV3 (ovarian carcinoma) (Buick et al., 1985b) were grown in complete DMEM (Corning) supplemented with 10% FBS (HyClone). Polyploid B16 x MEF fused clonal lines (Searles et al., 2018) and were kindly provided by Dr. Jack Bui (Dept. of Pathology, UCSD), and grown in complete RPMI media (Corning). All cells were maintained at 37 ◻C with 5% CO_2_ and were mycoplasma free as determined a PCR assay (Southern Biotech).

### BMDM generation in culture

Bone marrow derived macrophages (BMDM) were generated by isolating the femur and tibia of C57Bl/6 mice (8-12 weeks old) and flushing out the bone marrow using cold, unsupplemented RPMI growth medium (Corning) using a 27-gauge needle and syringe. Red cells were lysed using ACK Lysis buffer (Bio Whittaker). Macrophage differentiation from bone marrow cells was obtained by culture in standard growth medium supplemented with m-CSF (R&D Systems) at 30 ng/ml for 7 days.

### RNA isolation and cDNA synthesis

RNA was harvested from cells using Nucleopsin II Kit (Machery-Nagel). Concentration and purity of RNA were quantified the NanoDrop (ND-1000) spectrophotometer (Thermo Scientific) and analyzed with NanoDrop Software v3.8.0. RNA was normalized between conditions and cDNA generated using the High Capacity cDNA Synthesis kit (Life Technologies).

### XBP1 splicing assay

cDNA was subjected to the Xbp1 splicing assay as a surrogate outcome measure for ER stress. Primers were developed flanking the region of Xbp1 excised following UPR activation: Forward-5′-AGGGGAATGAAGTGAGGCCA-3′, Reverse - 5′-TGTGGTCAAAACGAATTAGT-3’. PCR was run on a Thermocycler (Thermo Scientific) using under the following conditions: 30 sec at 94°C, 40 sec at 55°C, 30 sec at 72°C for 35 cycles and 5min at 72°C. PCR products were run overnight on a 3% agarose gel at 30V for separation. Unspliced Xbp1 appears as the “upper band” at 358bp, while the spliced isoform appears as the “lower band” at 332bp. Data analysis and quantification of Xbp1 splicing was performed using ImageJ software.

### RT-qPCR

cDNA was subjected to RT-qPCR using an ABI 7300 Real-Time PCR system and TaqMan reagents for 50 cycles under universal cycling conditions. Cycling conditions followed manufacturer’s specifications (KAPA Biosystems). Target gene expression was normalized to β-actin and relative expression determined by using the -ΔΔCt relative quantification method. Primers for RT-qPCR were purchased from Life Technologies: Arg1 (Mm00475988_m1) and Il6 (Mm99999064_m1).

