## SUpplementray Material. Figures 1-14 and Tables 1-5 for "The unfolded protein response links tumor aneuploidy to local immune dysregulation": Supplementary Figures&Tables 11.09.20.pdf

**This PDF file includes:**

Figures S1 to S14

Tables S1 to S5

**Fig. S1**

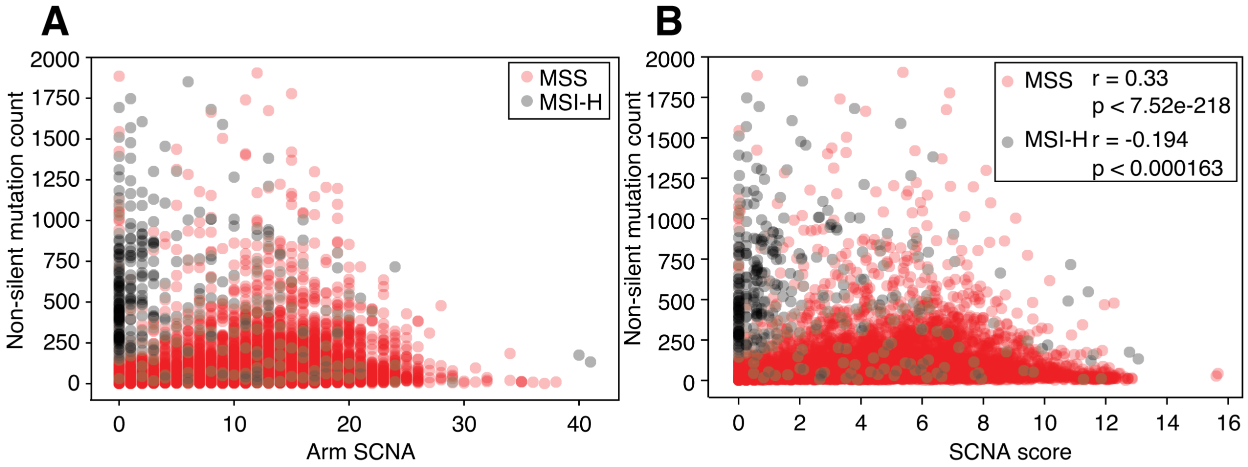


**Figure S1**. Pan-cancer analysis of somatic copy number alteration correlation with non-silent mutation burden

(A) Scatter plot of arm level somatic copy-number alteration count (x axis) versus non-silent mutation count (y axis). Red dots represent microsatellite stable (MSS) samples (N = 8536) and black dots represent microsatellite instability high (MSI-H) samples (N = 373).

(B) Scatter plot of tumor SCNA score (x axis) versus non-silent mutation count (y axis). Black dots represent MSI-H samples (N = 373) and red dots represent MSS samples (N = 8536). Spearman correlation for MSS samples is 0.33 ($p < 7.52e-218$). Spearman correlation for MSI-H samples is -0.194 ($p < 0.000163$).

**Fig. S2**

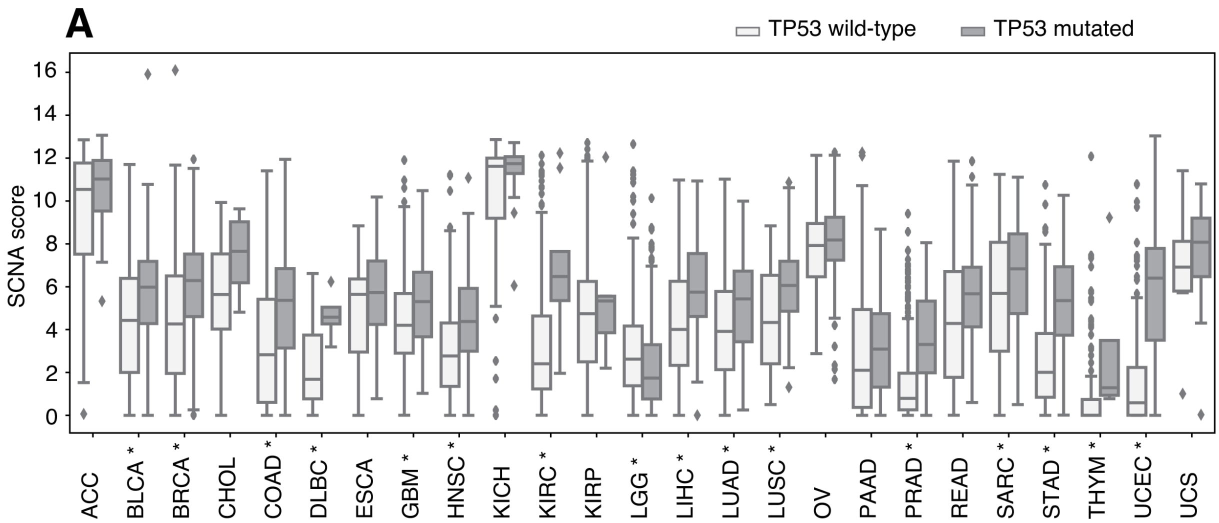
A
TP53 wild-type
TP53 mutated
SCNA score
16
14
12
10
8
6
4
2
0
ACC
BLCA *
BRCA *
CHOL
COAD *
DLBC *
ESCA
GBM *
HNSC *
KICH
KIRC *
KIRP
LGG *
LIHC *
LUAD *
LUSC *
OV
PAAD
PRAD *
READ
SARC *
STAD *
THYM *
UCEC *
UCS

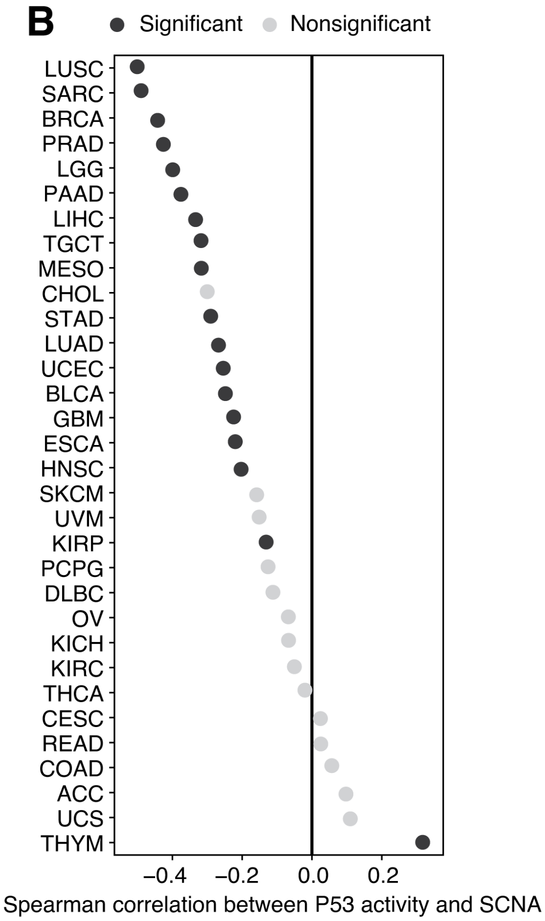
B
Significant
Nonsignificant
LUSC
SARC
BRCA
PRAD
LGG
PAAD
LIHC
TGCT
MESO
CHOL
STAD
LUAD
UCEC
BLCA
GBM
ESCA
HNSC
SKCM
UVM
KIRP
PCPG
DLBC
OV
KICH
KIRC
THCA
CESC
READ
COAD
ACC
UCS
THYM
−0.4
−0.2
0.0
0.2
Spearman correlation between P53 activity and SCNA

**Figure S2**. p53 activity correlates with SCNA score.

(A) Boxplots showing SCNA score distribution differences between TP53 mutated samples and TP53 wild-type samples across 25 tumor types with TP53 mutations.

(B) Spearman correlation between p53 activity and SCNA score across 32 tumor types. Black dots represent statistically significant values (FDR < 0.05) and gray dots represents non-significant values (FDR > 0.05).

**Fig. S3**

**A**

| | Chr | Arm | Focal | SCNA |
|---|---|---|---|---|
| Chr | 1.000 | 0.548 | 0.612 | 0.860 |
| Arm | 0.548 | 1.000 | 0.627 | 0.866 |
| Focal | 0.612 | 0.627 | 1.000 | 0.735 |
| SCNA | 0.860 | 0.866 | 0.735 | 1.000 |

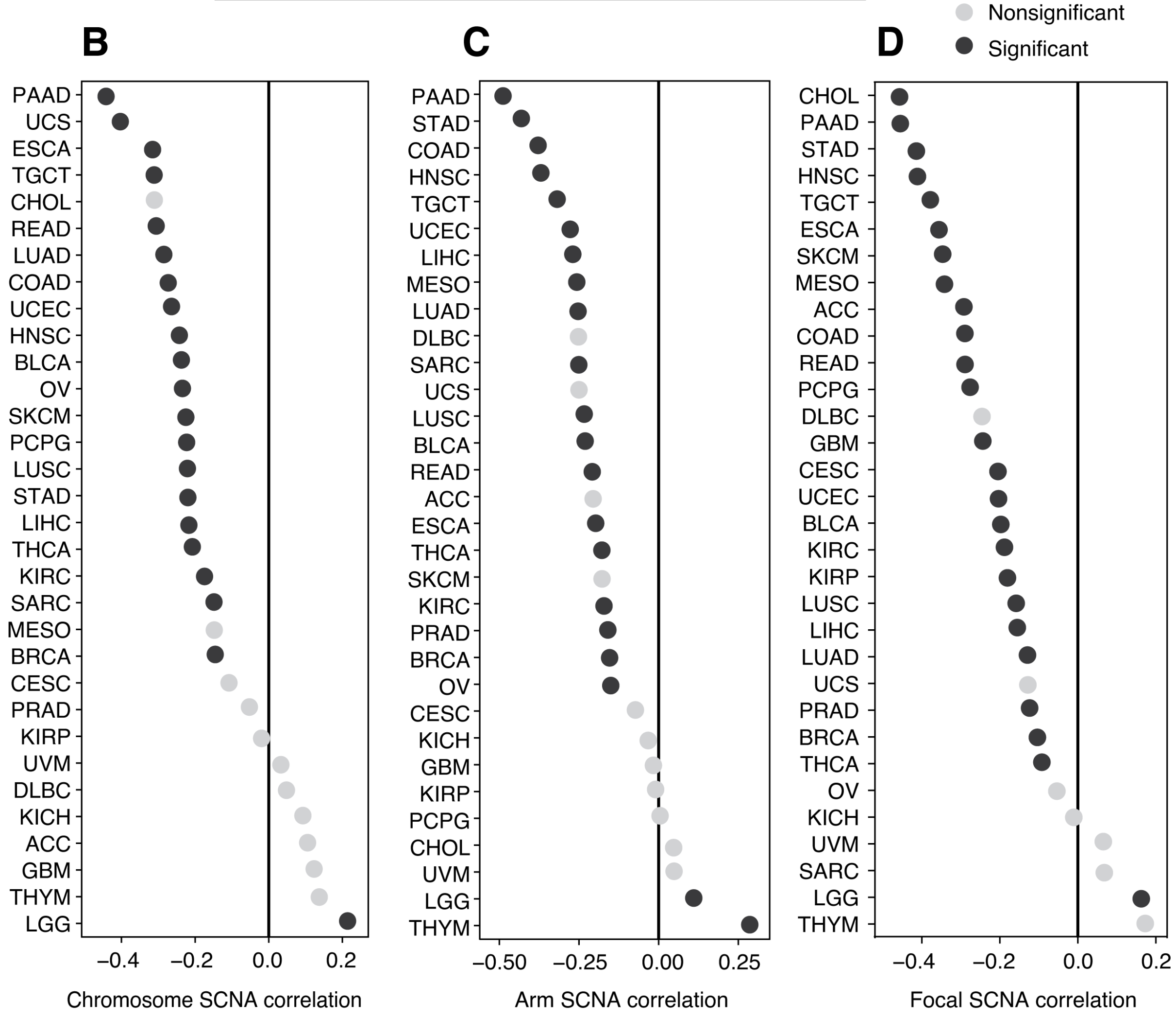

**Figure S3**. Three categories of SCNA show inverse correlation with CYT score across tumor types

(A) Pairwise Spearman correlation coefficients show that the 3 categories of SCNA and the combined single SCNA score are positively correlated pancancer.

(B) Chromosome copy-number alteration count correlation with CYT score across 32 tumor types.

(C) Chromosome arm copy-number alteration count correlation with CYT score across 32 tumor types.

(D) Focal copy-number alteration count correlation with CYT score across 32 tumor types.

**Fig. S4**

**A**

| | coeff | p-value |
|---|---|---|
| chr | -0.145 | 2.76e-30 |
| arm | -0.198 | 1.70e-50 |
| focal | -0.070 | 1.65e-09 |

**B**

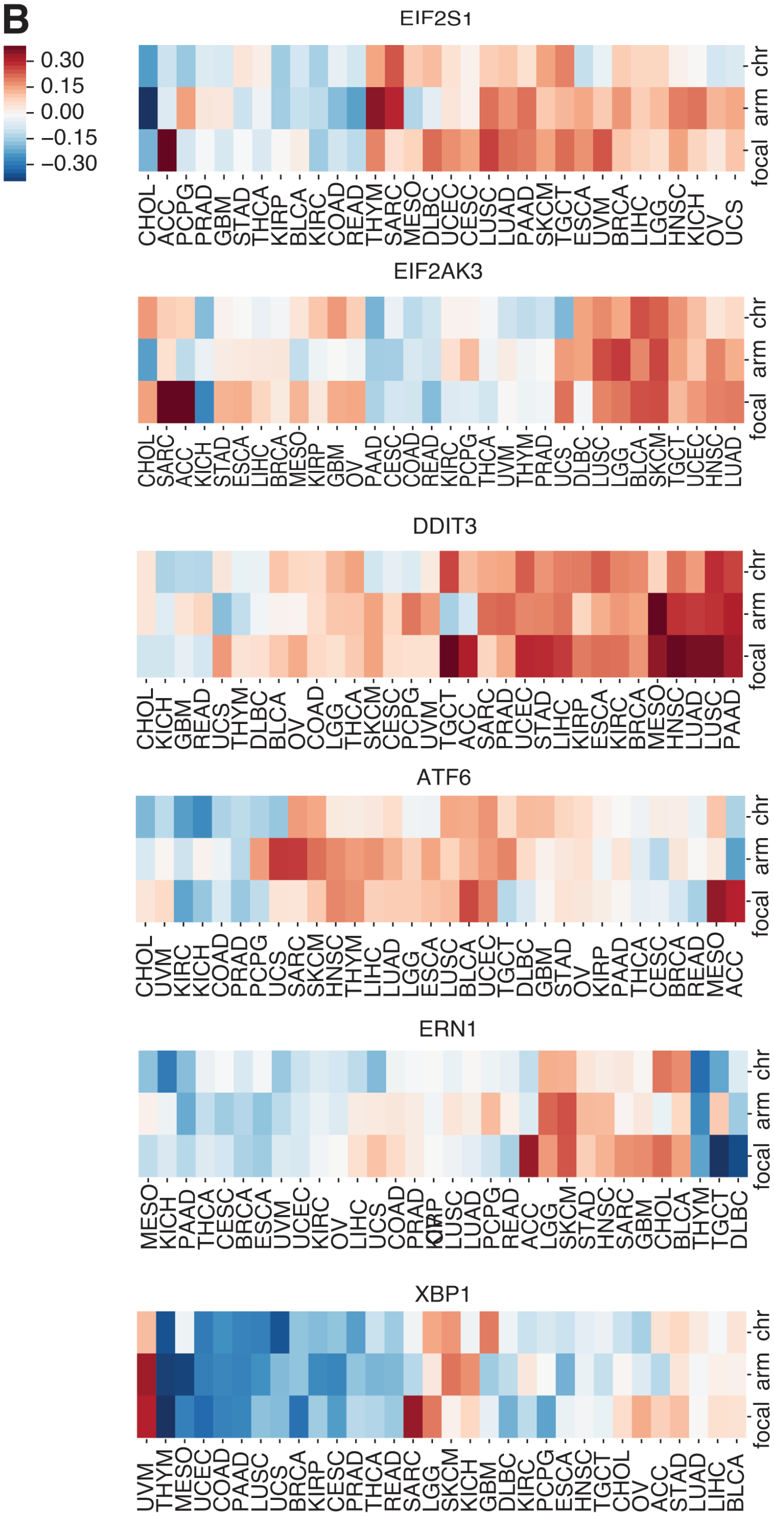

**Figure S4**. Three categories of SCNA event show similar trends in correlation with CYT score UPR gene expression

(A) Coefficients and p-values of an ordinary least square (OLS) linear model using 3 categories of SCNA to predict CYT score, with tumor type as a covariate.

(B) Heatmaps showing correlation between the 3 categories of SCNA event (rows) and UPR parental genes for 32 tumor types (columns). Color intensity indicates magnitude of Spearman correlation, with red indicating positive and blue indicating negative correlation respectively.

**Fig. S5**

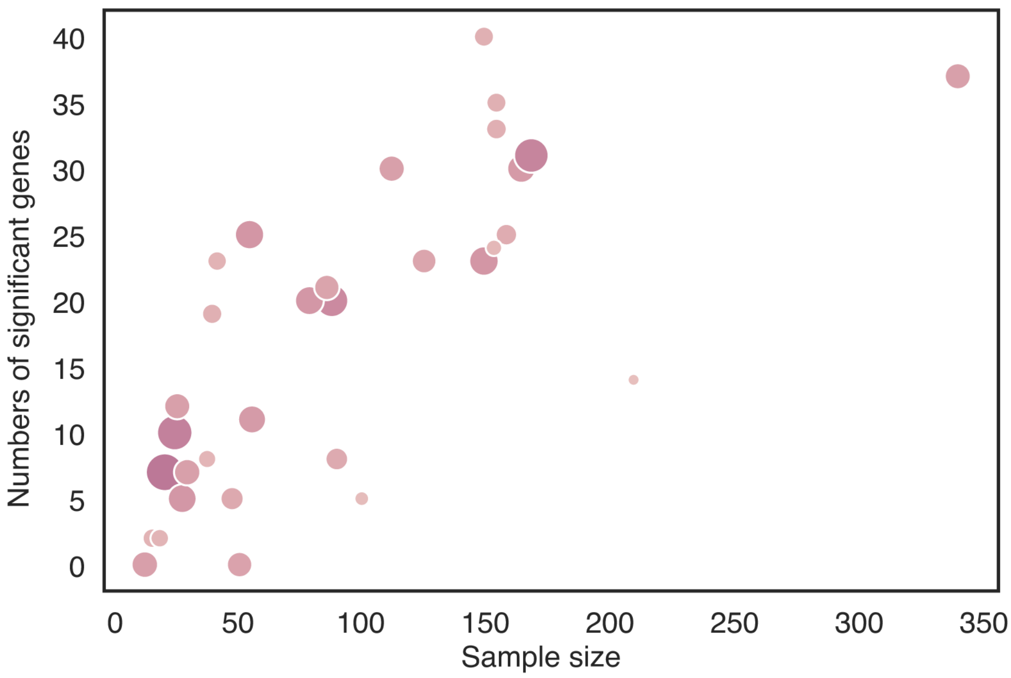


**Figure S5**. Sample size limits the number of significant differentially expressed genes detected between SCNA$^{high}$ and SCNA$^{low}$ tumors

A scatter plot showing the number of samples (x axis) versus the number of UPR genes with significant differential expression after multiple hypothesis testing (y axis). Each dot represents a tumor type in TCGA. The size and color intensity of the dots depicts the median variance across UPR genes for a tumor type.

**Fig. S6**

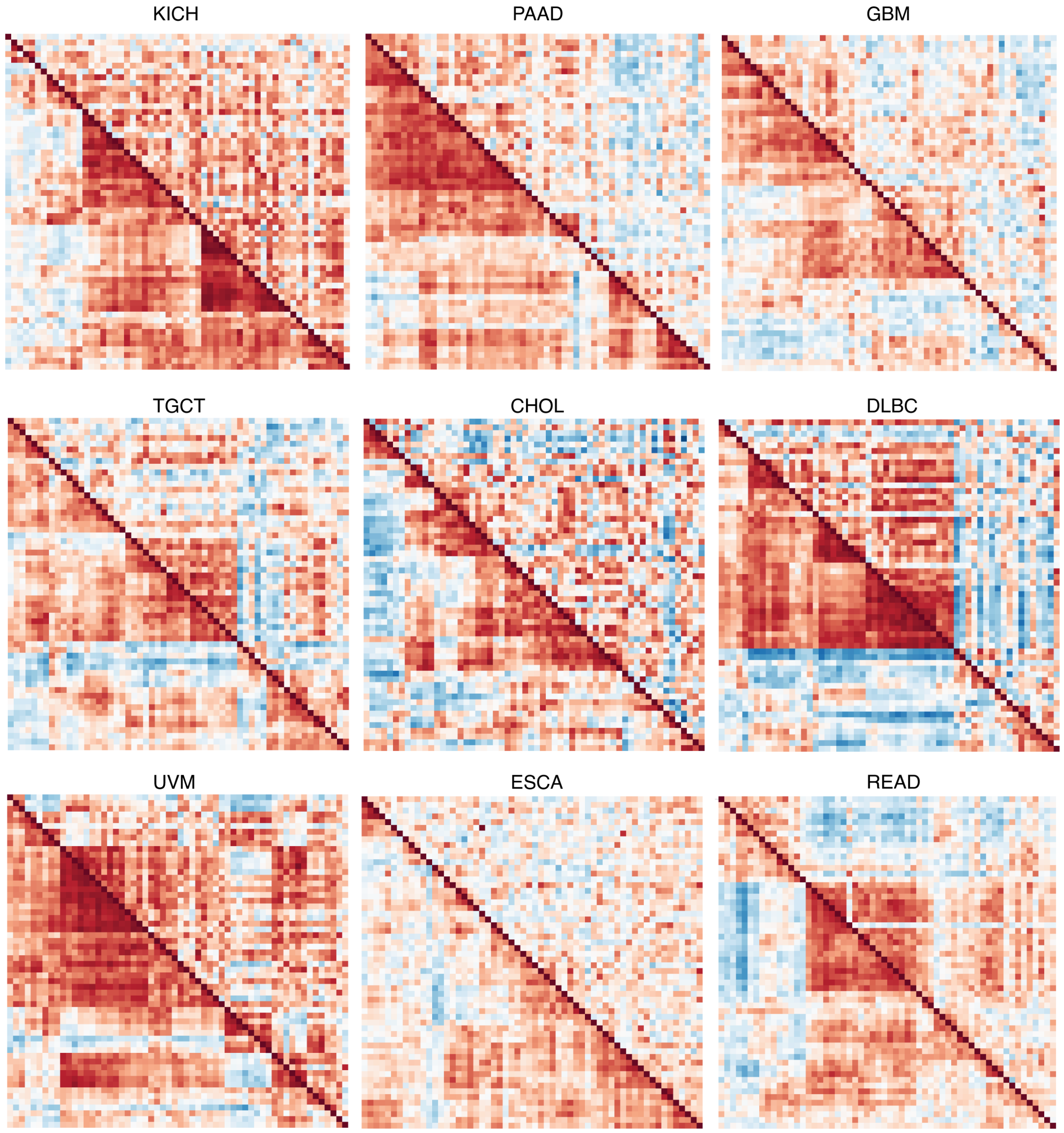


**Figure S6**. Heatmaps showing UPR gene co-expression patterns for high and low SCNA groups for 9 tumor types. Bottom left triangles represent the co-expression of UPR genes in the SCNA$^{low}$ group, and top right triangles represent the co-expression of UPR genes in the SCNA$^{high}$ group.

### Fig. S7

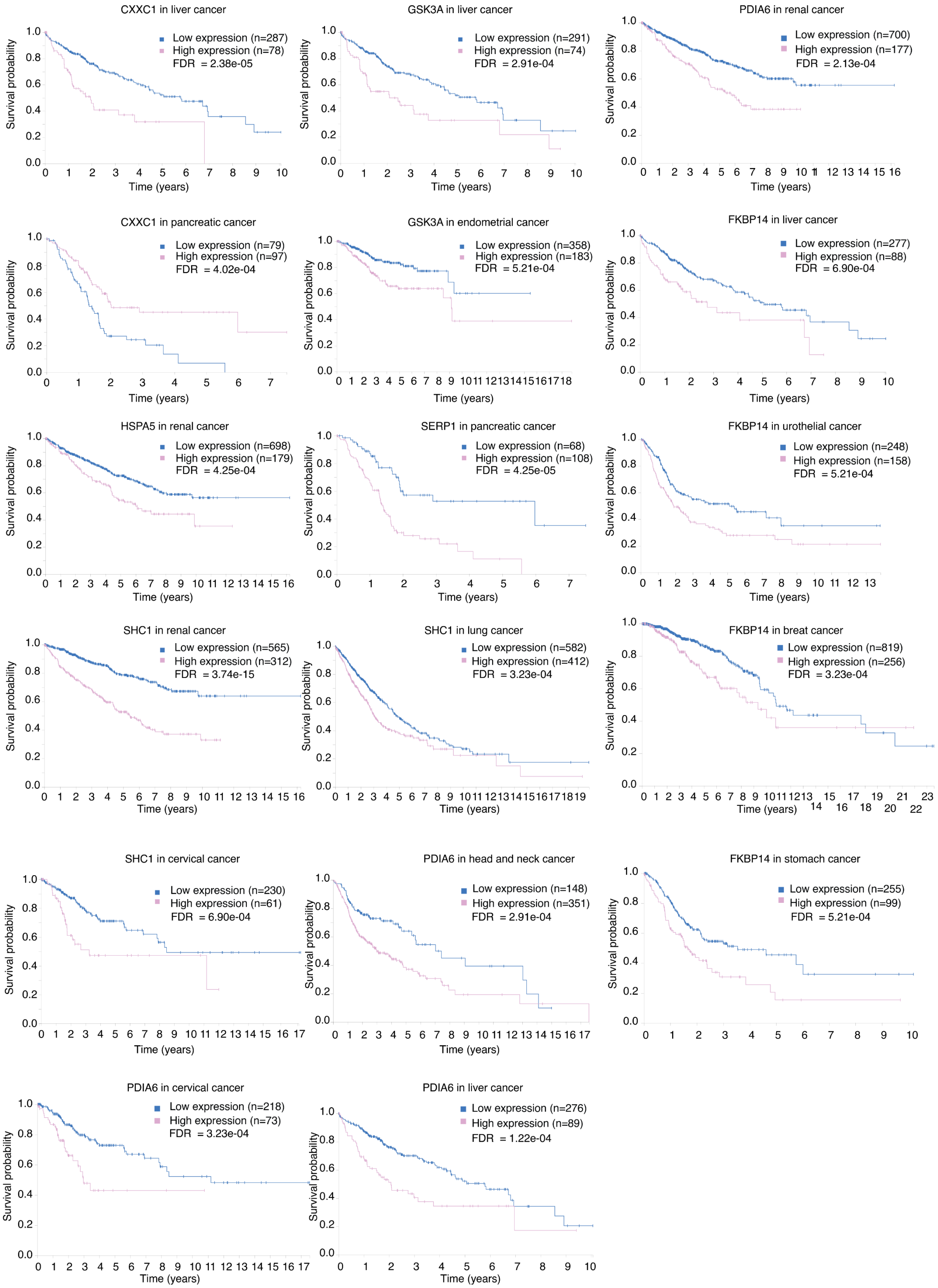
CXXC1 in liver cancer
Low expression (n=287)
High expression (n=78)
FDR = 2.38e-05
GSK3A in liver cancer
Low expression (n=291)
High expression (n=74)
FDR = 2.91e-04
PDIA6 in renal cancer
Low expression (n=700)
High expression (n=177)
FDR = 2.13e-04
CXXC1 in pancreatic cancer
Low expression (n=79)
High expression (n=97)
FDR = 4.02e-04
GSK3A in endometrial cancer
Low expression (n=358)
High expression (n=183)
FDR = 5.21e-04
FKBP14 in liver cancer
Low expression (n=277)
High expression (n=88)
FDR = 6.90e-04
HSPA5 in renal cancer
Low expression (n=698)
High expression (n=179)
FDR = 4.25e-04
SERP1 in pancreatic cancer
Low expression (n=68)
High expression (n=108)
FDR = 4.25e-05
FKBP14 in urothelial cancer
Low expression (n=248)
High expression (n=158)
FDR = 5.21e-04
SHC1 in renal cancer
Low expression (n=565)
High expression (n=312)
FDR = 3.74e-15
SHC1 in lung cancer
Low expression (n=582)
High expression (n=412)
FDR = 3.23e-04
FKBP14 in breat cancer
Low expression (n=819)
High expression (n=256)
FDR = 3.23e-04
SHC1 in cervical cancer
Low expression (n=230)
High expression (n=61)
FDR = 6.90e-04
PDIA6 in head and neck cancer
Low expression (n=148)
High expression (n=351)
FDR = 2.91e-04
FKBP14 in stomach cancer
Low expression (n=255)
High expression (n=99)
FDR = 5.21e-04
PDIA6 in cervical cancer
Low expression (n=218)
High expression (n=73)
FDR = 3.23e-04
PDIA6 in liver cancer
Low expression (n=276)
High expression (n=89)
FDR = 1.22e-04
Survival probability
Time (years)

**Figure S7**. Poor prognosis linked to genes with loss of coordination under SCNA$^{high}$ conditions. Genes with reduced coordination in expression with three or more partners (*CXXC1, GSK3A, HSPA5, SERP1, SHC1, PDIA6* and *FKBP14)* were input to the HUMAN PROTEIN ATLAS (version 19.3) at http://www.proteinatlas.org, DOI: 10.1126/science.aan2507. This site performs a Kaplan-Meier analysis in 17 tumor types from TCGA, stratifying samples into high or low expression level based on predetermined FPKM cutoffs (55). Plots are returned only if the log-rank test was significant. P values and multiple testing adjusted p values are shown with each plot. In general, higher expression of UPR genes was associated with poorer outcome.

#### Fig S8

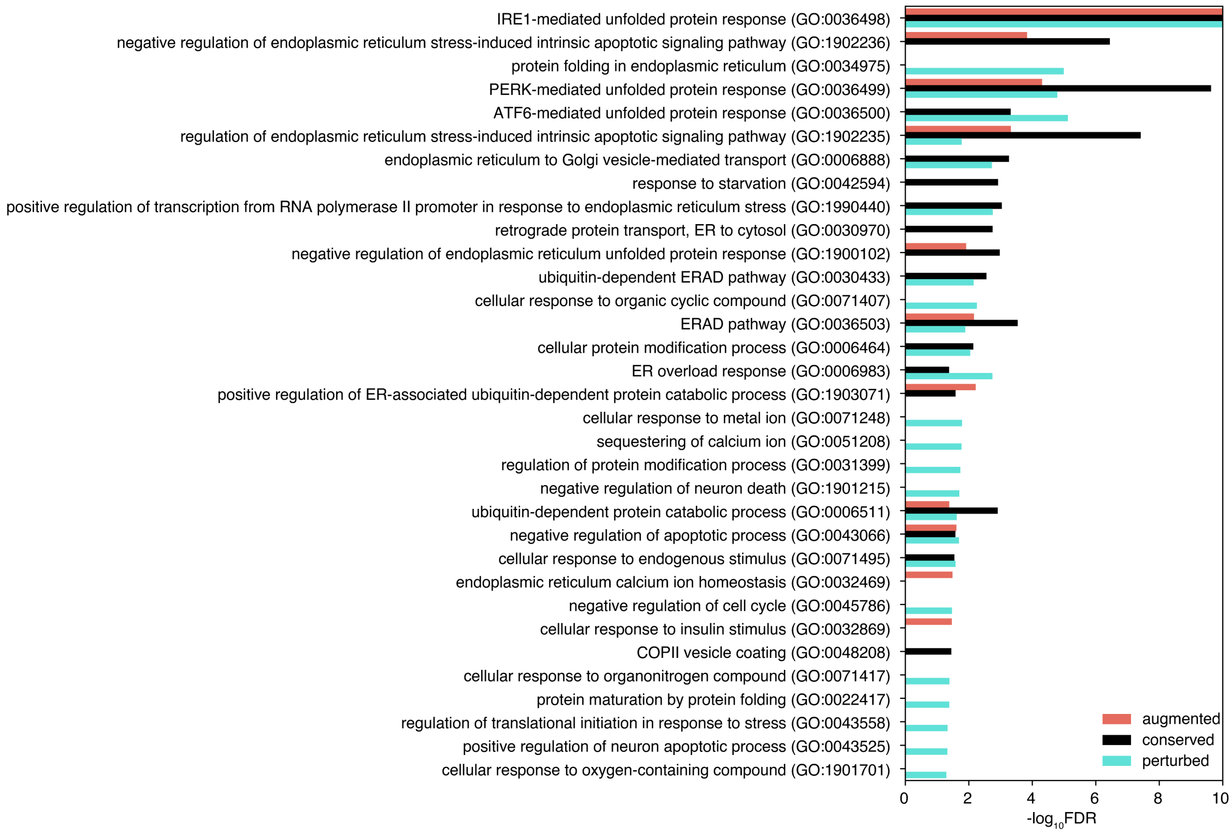


**Figure S8**. Gene Ontology Analysis for perturbed, preserved and augmented gene pairs, showing only parental Gene Ontology terms. Negative log10 adjusted p-values of GO terms enriched for preserved, augmented or perturbed genes ordered by median across categories, focusing on parental terms from the GO hierarchy tree. The complete list is available in Table S3.

**Fig. S9**

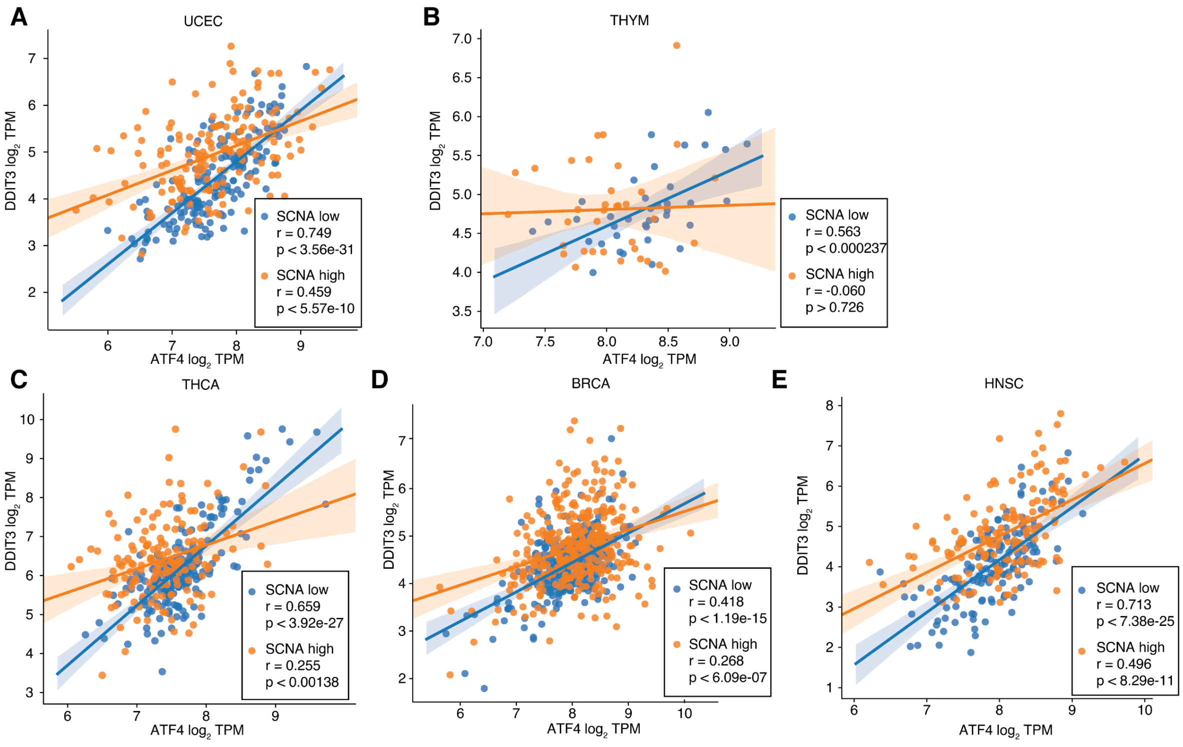


**Figure S9**. ATF4 and DDIT3 gene loss of coordination in four tumor types. Scatter plots showing the $\log_2$ TPM expression level of ATF4 (x axis) and DDIT3 (y axis) in $SCNA^{low}$ (blue) and $SCNA^{high}$ (orange) conditions for tumors from 4 different tumor types: (A) UCEC - Endometrial cancer; (B) THYM – Thymoma; (C) THCA – Thyroid cancer; (D) BRCA – Breast Adenocarcinoma; (E) HNSC – Head and neck squamous cell carcinoma. Regression lines indicate the strength of correlation between the expression of the two genes. Spearman correlation coefficients and associated p values are provided.

**Fig. S10**

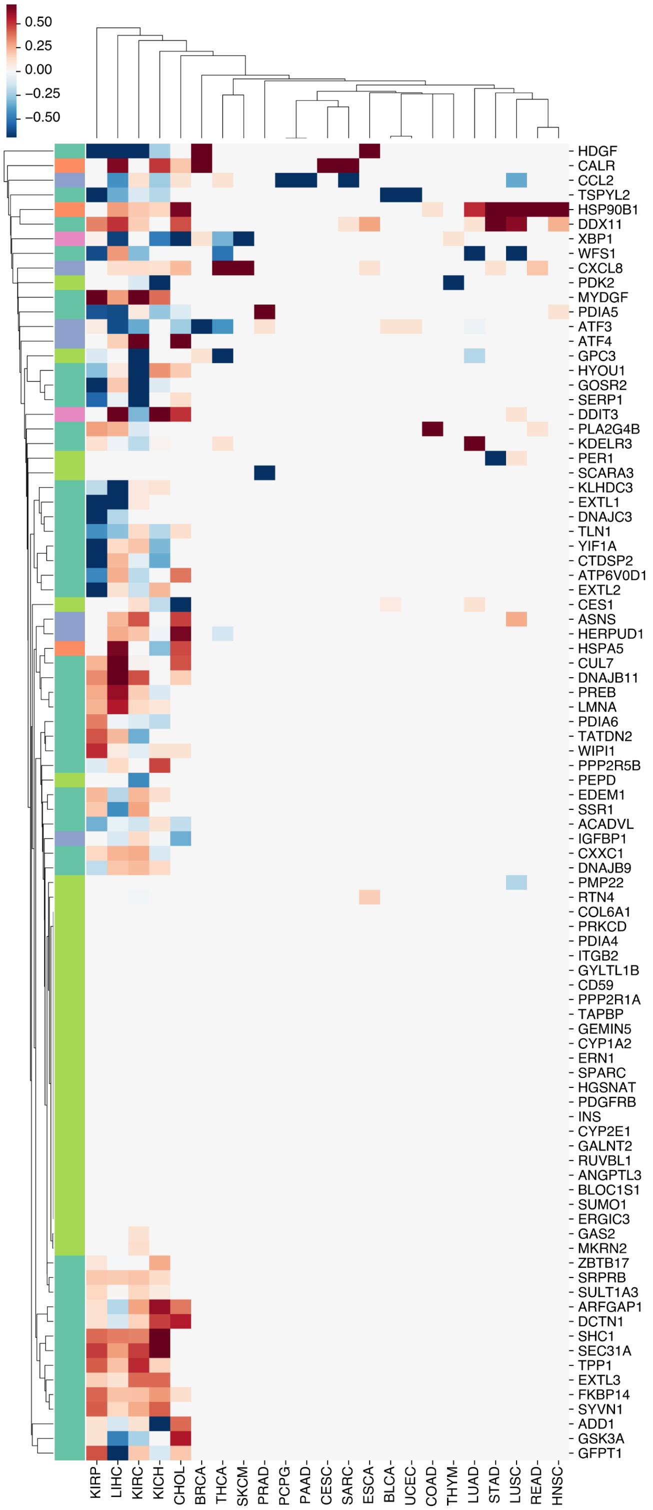

0.50
0.25
0.00
−0.25
−0.50
HDGF
CALR
CCL2
TSPYL2
HSP90B1
DDX11
XBP1
WFS1
CXCL8
PDK2
MYDGF
PDIA5
ATF3
ATF4
GPC3
HYOU1
GOSR2
SERP1
DDIT3
PLA2G4B
KDELR3
PER1
SCARA3
KLHDC3
EXTL1
DNAJC3
TLN1
YIF1A
CTDSP2
ATP6V0D1
EXTL2
CES1
ASNS
HERPUD1
HSPA5
CUL7
DNAJB11
PREB
LMNA
PDIA6
TATDN2
WIPI1
PPP2R5B
PEPD
EDEM1
SSR1
ACADVL
IGFBP1
CXXC1
DNAJB9
PMP22
RTN4
COL6A1
PRKCD
PDIA4
ITGB2
GYLTL1B
CD59
PPP2R1A
TAPBP
GEMIN5
CYP1A2
ERN1
SPARC
HGSNAT
PDGFRB
INS
CYP2E1
GALNT2
RUVBL1
ANGPTL3
BLOC1S1
SUMO1
ERGIC3
GAS2
MKRN2
ZBTB17
SRPRB
SULT1A3
ARFGAP1
DCTN1
SHC1
SEC31A
TPP1
EXTL3
FKBP14
SYVN1
ADD1
GSK3A
GFPT1
KIRP
LIHC
KIRC
KICH
CHOL
BRCA
THCA
SKCM
PRAD
PCPG
PAAD
CESC
SARC
ESCA
BLCA
UCEC
COAD
THYM
LUAD
STAD
LUSC
READ
HNSC

**Figure S10**. Genes selected to represent UPR pathway activity by Lasso regression for 23 tumor types. A heatmap showing genes selected by Lasso regression models for three UPR pathways and 23 tumor types during pathway score quantification. Each column represents a tumor type and each row represents a gene. The left color bar indicates gene membership in UPR branch pathways: IRE1 – green; PERK – blue; ATF6 – orange; RIDD – light green; multiple pathways – purple. Each cell in the heatmap is colored according to the magnitude of the coefficient of the gene in the lasso regression model within each tumor type, with red depicting positive coefficients and blue depicting negative coefficients. To achieve a better visualization, coefficients were scaled within each tumor type to fall in the -1 to 1 range and thus represent the relative strength of gene contribution for different tumor types.

**Fig. S11**

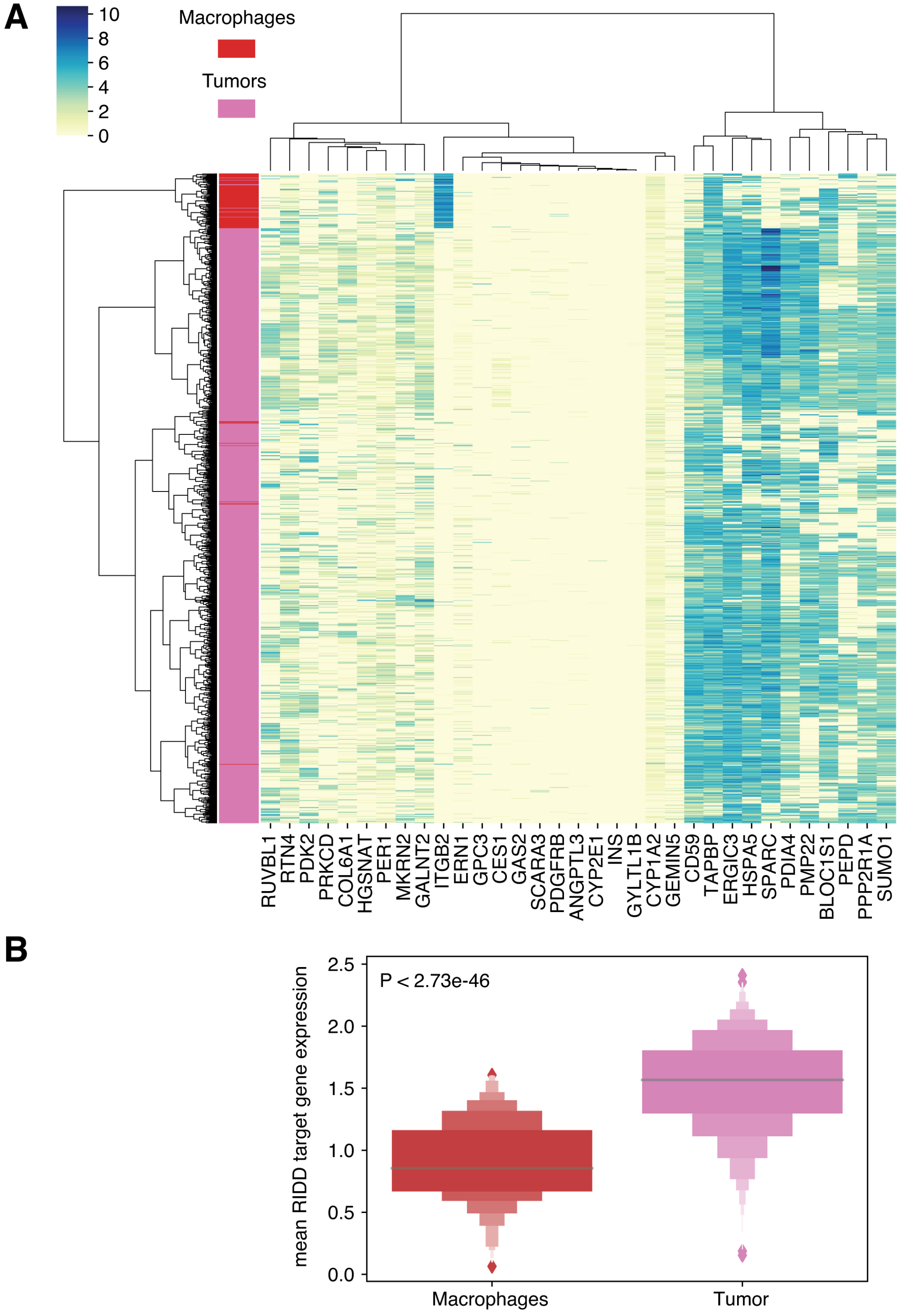
A
10
8
6
4
2
0
Macrophages
Tumors
RUVBL1
RTN4
PDK2
PRKCD
COL6A1
HGSNAT
PER1
MKRN2
GALNT2
ITGB2
ERN1
GPC3
CES1
GAS2
SCARA3
PDGFRB
ANGPTL3
CYP2E1
INS
GYLTL1B
CYP1A2
GEMIN5
CD59
TAPBP
ERGIC3
HSPA5
SPARC
PDIA4
PMP22
BLOC1S1
PEPD
PPP2R1A
SUMO1
B
P < 2.73e-46
mean RIDD target gene expression
2.5
2.0
1.5
1.0
0.5
0.0
Macrophages
Tumor

**Figure S11**. Analysis of RIDD target gene expression in single-cell data showed reduced expression in tumor-associated macrophages compared to tumor cells.

(A) A heatmap showing the unsupervised clustering of 1257 tumor cells and 119 macrophages (rows) according to expression in XX of 33 RIDD target genes (columns). The left sidebar indicates cell type: red – macrophages; pink – tumor cells.

(B) Boxplots showing the distribution of mean RIDD target gene expression in macrophages (blue) and tumor cells (orange). ITGB2 and TAPBP were excluded as their behavior is counter to regulation by RIDD.

**Fig. S12**

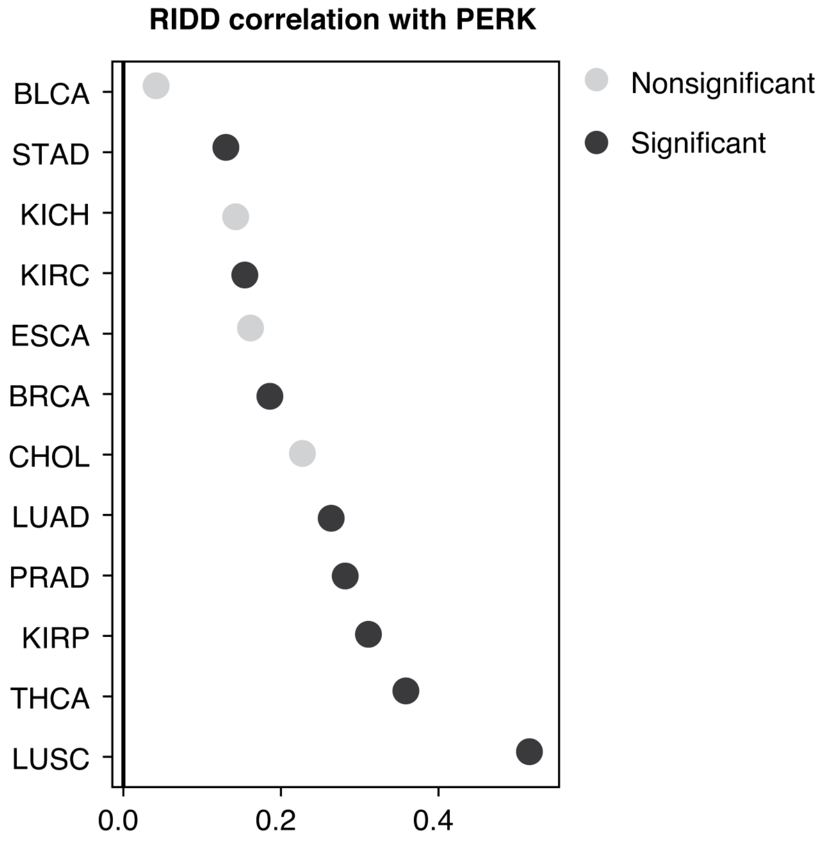


**Figure S12**. Spearman correlation coefficients linking PERK pathway score and RIDD activity score for 13 tumor types for which both pathway scores could be calculated. Black color indicates correlations that were statistically significant after multiple hypothesis testing correction using the Benjamini Hochberg procedure (FDR < 0.05). Gray dots represent non-significant cases.

**Fig. S13.**

Chromosome 11, DLD1

Unstimulated

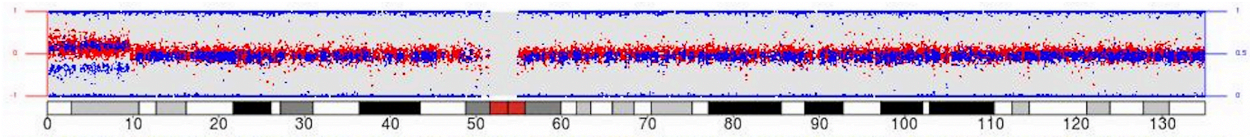


+Rv (14 days)

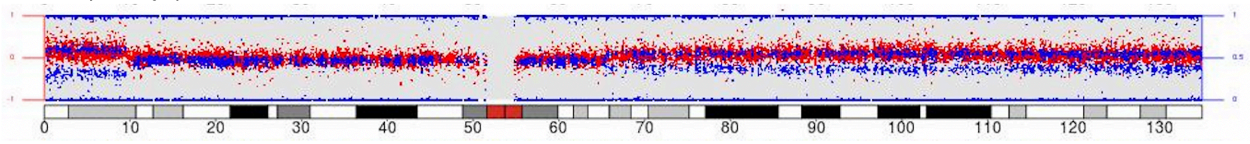


Chromosome 20, DLD1

Unstimulated

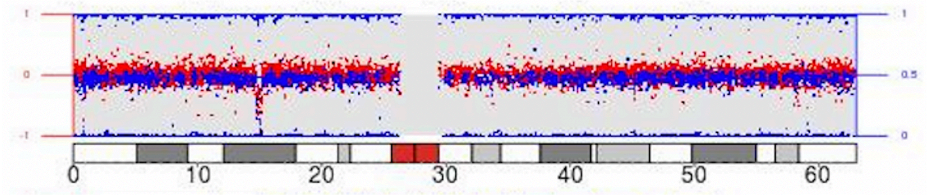


+Rv (14 days)

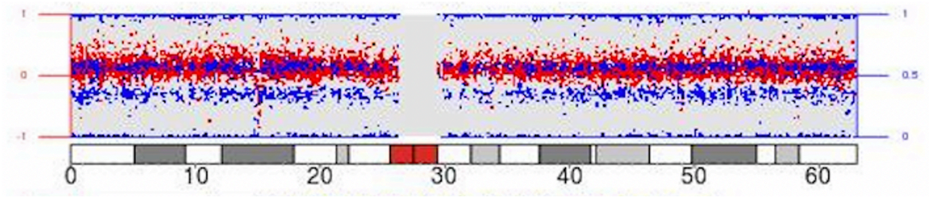


**Figure S13**. Digital Karyotyping for Detecting Chromosome Alterations. Mean log R ratio (LRR, in red) and B-allele frequency (BAF, in blue). A. AAB/ABB trisomy specific BAF signal (the two blue horizontal bands) at 11p is present both before and after stimulation, while only stimulated cells show AAB/ABB trisomy specific BAF signal for 11q. On chromosome 20, a small deletion is observed in both untreated and treated cells at 20p, while AAB/ABB trisomy for the entirety of chromosome 20 was observed only in treated cells.

**Fig. S14.**

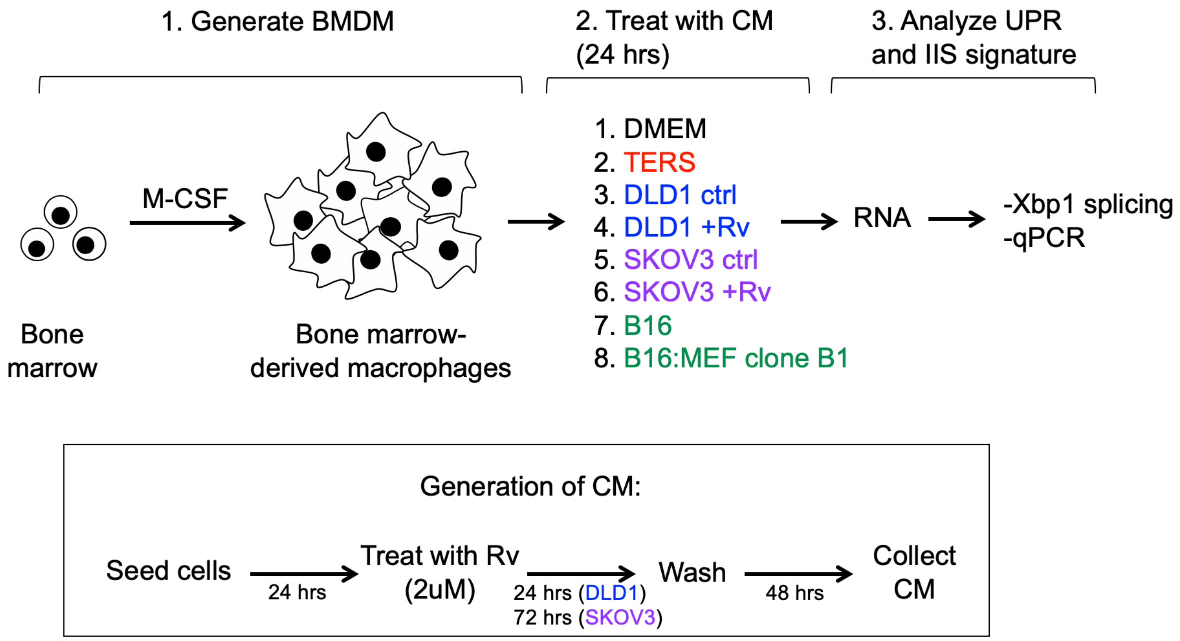


**Figure S14**. Schematic representation of the workflow for transmissible immune dysregulation experiments

### Supplementary Tables

**Table S1.** Genes associated with four REACTOME pathways used to represent IRE1α, PERK, ATF6 and XBP1 pathways.

| | PERK (R-HSA | ATF6 (R-HSA | IRE1alpha (R | XBP1S (R-HSA-381038.2) |
|---|---|---|---|---|
| 0 | ASNS | DDIT3 | ADD1 | ADD1 |
| 1 | IGFBP1 | HSP90B1 | YIF1A | YIF1A |
| 2 | DDIT3 | CALR | TLN1 | TLN1 |
| 3 | CCL2 | HSPA5 | TPP1 | TPP1 |
| 4 | HERPUD1 | XBP1 | DNAJB11 | DNAJB11 |
| 5 | ATF3 | | SYVN1 | SYVN1 |
| 6 | ATF4 | | EXTL3 | EXTL3 |
| 7 | CXCL8 | | PREB | PREB |
| 8 | | | EXTL1 | EXTL1 |
| 9 | | | TATDN2 | TATDN2 |
| 10 | | | DCTN1 | DCTN1 |
| 11 | | | DNAJC3 | DNAJC3 |
| 12 | | | CXXC1 | CXXC1 |
| 13 | | | GFPT1 | GFPT1 |
| 14 | | | GOSR2 | GOSR2 |
| 15 | | | EDEM1 | EDEM1 |
| 16 | | | SSR1 | SSR1 |
| 17 | | | SERP1 | SERP1 |
| 18 | | | PPP2R5B | PPP2R5B |
| 19 | | | WIPI1 | WIPI1 |
| 20 | | | SULT1A3 | SULT1A3 |
| 21 | | | WFS1 | WFS1 |
| 22 | | | DDX11 | DDX11 |
| 23 | | | GSK3A | GSK3A |
| 24 | | | HDGF | HDGF |
| 25 | | | KLHDC3 | KLHDC3 |
| 26 | | | FKBP14 | FKBP14 |
| 27 | | | SRPRB | SRPRB |
| 28 | | | DNAJB9 | DNAJB9 |
| 29 | | | TSPYL2 | TSPYL2 |
| 30 | | | ZBTB17 | ZBTB17 |
| 31 | | | PDIA5 | PDIA5 |
| 32 | | | CUL7 | CUL7 |
| 33 | | | KDELR3 | KDELR3 |
| 34 | | | SEC31A | SEC31A |
| 35 | | | ACADVL | ACADVL |
| 36 | | | SHC1 | SHC1 |
| 37 | | | PLA2G4B | PLA2G4B |
| 38 | | | EXTL2 | EXTL2 |
| 39 | | | LMNA | LMNA |
| 40 | | | ARFGAP1 | ARFGAP1 |
| 41 | | | HYOU1 | HYOU1 |
| 42 | | | CTDSP2 | CTDSP2 |
| 43 | | | PDIA6 | PDIA6 |
| 44 | | | ATP6V0D1 | ATP6V0D1 |
| 45 | | | XBP1 | MYDGF |
| 46 | | | MYDGF | |

**Table S2.** A list of gene pairs with preserved, augmented or perturbed co-expression patterns used to construct the network in main Figure 4D.

| | source | target | condition |
|---|---|---|---|
| 0 | FKBP14 | GOSR2 | perturbed |
| 1 | ACADVL | KDELR3 | augmented |
| 2 | EDEM1 | TPP1 | perturbed |
| 3 | DCTN1 | KLHDC3 | perturbed |
| 4 | SRPRB | ZBTB17 | augmented |
| 5 | HDGF | TATDN2 | perturbed |
| 6 | ATP6V0D1 | GSK3A | perturbed |
| 7 | EXTL2 | SHC1 | perturbed |
| 8 | GOSR2 | SHC1 | perturbed |
| 9 | HYOU1 | PDIA5 | perturbed |
| 10 | SERP1 | ZBTB17 | augmented |
| 11 | HYOU1 | SERP1 | perturbed |
| 12 | GSK3A | PPP2R5B | perturbed |
| 13 | ATP6V0D1 | HSP90B1 | perturbed |
| 14 | ATP6V0D1 | PDIA6 | perturbed |
| 15 | CXCL8 | HDGF | perturbed |
| 16 | KDELR3 | TPP1 | perturbed |
| 17 | CTDSP2 | FKBP14 | perturbed |
| 18 | ACADVL | CXXC1 | perturbed |
| 19 | EDEM1 | GOSR2 | perturbed |
| 20 | TLN1 | WFS1 | perturbed |
| 21 | KLHDC3 | PPP2R5B | perturbed |
| 22 | EXTL2 | SEC31A | perturbed |
| 23 | DCTN1 | GSK3A | perturbed |
| 24 | SRPRB | SSR1 | perturbed |
| 25 | HSPA5 | SERP1 | perturbed |
| 26 | CALR | HSPA5 | perturbed |
| 27 | HSPA5 | HYOU1 | perturbed |
| 28 | SYVN1 | YIF1A | augmented |
| 29 | SHC1 | TPP1 | perturbed |
| 30 | CALR | EXTL3 | perturbed |
| 31 | CXXC1 | SULT1A3 | perturbed |
| 32 | DNAJC3 | SERP1 | perturbed |
| 33 | SERP1 | SSR1 | perturbed |
| 34 | CXXC1 | SSR1 | augmented |
| 35 | ATF4 | CXXC1 | perturbed |
| 36 | HERPUD1 | HSPA5 | augmented |
| 37 | ATF4 | SULT1A3 | perturbed |
| 38 | ATF4 | PPP2R5B | perturbed |
| 39 | FKBP14 | SYVN1 | preserved |
| 40 | FKBP14 | EXTL1 | preserved |
| 41 | FKBP14 | SULT1A3 | preserved |

**Table S3.** Go Ontology Analysis. The complete list of GO terms enriched for preserved, augmented or perturbed genes by categories (preserved, augmented or perturbed). Negative $\log_{10}$ adjusted p-values are provided.

| | GO biologica | negative log: | condition |
|---|---|---|---|
| 0 | positive regu | 2.22 | augmented |
| 1 | integrated st | 4.28 | augmented |
| 2 | PERK-mediat | 4.31 | augmented |
| 3 | regulation of | 2.08 | augmented |
| 4 | positive regu | 4.01 | augmented |
| 5 | negative regu | 1.92 | augmented |
| 6 | negative regu | 3.84 | augmented |
| 7 | IRE1-mediat | 14.79 | augmented |
| 8 | ER-nucleus s | 7.62 | augmented |
| 9 | endoplasmic | 20.85 | augmented |
| 10 | regulation of | 3.38 | augmented |
| 11 | regulation of | 3.32 | augmented |
| 12 | cellular resp | 20.21 | augmented |
| 13 | endoplasmic | 1.49 | augmented |
| 14 | positive regu | 3.14 | augmented |
| 15 | cellular resp | 19.75 | augmented |
| 16 | regulation of | 1.42 | augmented |
| 17 | response to u | 19.35 | augmented |
| 18 | negative regu | 2.92 | augmented |
| 19 | response to t | 18.89 | augmented |
| 20 | regulation of | 3.97 | augmented |
| 21 | response to e | 17.49 | augmented |
| 22 | ERAD pathw | 2.16 | augmented |
| 23 | negative regu | 2.07 | augmented |
| 24 | positive regu | 1.94 | augmented |
| 25 | positive regu | 1.77 | augmented |
| 26 | positive regu | 1.61 | augmented |
| 27 | regulation of | 1.53 | augmented |
| 28 | cellular resp | 1.47 | augmented |
| 29 | cellular resp | 1.39 | augmented |
| 30 | regulation of | 1.39 | augmented |
| 31 | ubiquitin-dep | 1.38 | augmented |
| 32 | modification | 1.38 | augmented |
| 33 | modification | 1.36 | augmented |
| 34 | cellular resp | 1.32 | augmented |
| 35 | response to c | 3.84 | augmented |
| 36 | response to r | 3.67 | augmented |
| 37 | cellular resp | 8.49 | augmented |
| 38 | negative regu | 1.61 | augmented |
| 39 | cellular hom | 1.6 | augmented |
| 40 | negative regu | 1.61 | augmented |
| 41 | negative regu | 1.45 | augmented |

| | | | |
|---|---|---|---|
| 85 | vesicle targe | 2.25 | preserved |
| 86 | COPII-coated | 1.38 | preserved |
| 87 | regulation of | 3.66 | preserved |
| 88 | endoplasmic | 3.26 | preserved |
| 89 | response to : | 2.92 | preserved |
| 90 | negative reg | 1.94 | preserved |
| 91 | cellular resp | 32.61 | preserved |
| 92 | Golgi vesicle | 2.76 | preserved |
| 93 | proteasomal | 2.13 | preserved |
| 94 | regulation of | 2.72 | preserved |
| 95 | negative reg | 3 | preserved |
| 96 | proteasome- | 1.4 | preserved |
| 97 | cellular resp | 30.59 | preserved |
| 98 | ubiquitin-dep | 2.91 | preserved |
| 99 | modification | 2.9 | preserved |
| 100 | modification | 2.86 | preserved |
| 101 | proteolysis ir | 2.66 | preserved |
| 102 | cellular prote | 2.53 | preserved |
| 103 | cellular resp | 27.34 | preserved |
| 104 | response to r | 1.45 | preserved |
| 105 | regulation of | 2.85 | preserved |
| 106 | response to ( | 26.86 | preserved |
| 107 | protein catab | 2.27 | preserved |
| 108 | response to ( | 1.37 | preserved |
| 109 | response to ( | 3.45 | preserved |
| 110 | response to r | 3.16 | preserved |
| 111 | response to : | 24.04 | preserved |
| 112 | negative reg | 1.98 | preserved |
| 113 | negative reg | 1.58 | preserved |
| 114 | cellular macr | 1.55 | preserved |
| 115 | negative reg | 1.54 | preserved |
| 116 | negative reg | 2.63 | preserved |
| 117 | organonitrog | 1.84 | preserved |
| 118 | response to ( | 21.1 | preserved |
| 119 | negative reg | 2.26 | preserved |
| 120 | negative reg | 2.26 | preserved |
| 121 | cellular resp | 1.55 | preserved |
| 122 | signal transd | 19.38 | preserved |
| 123 | intracellular | 2.04 | preserved |
| 124 | establishmer | 1.9 | preserved |
| 125 | signaling (GC | 18.28 | preserved |
| 126 | cell commun | 18.05 | preserved |
| 127 | cellular catal | 2.12 | preserved |

| | | | |
|---|---|---|---|
| 128 | protein trans | 1.46 | preserved |
| 129 | regulation of | 1.68 | preserved |
| 130 | peptide trans | 1.42 | preserved |
| 131 | regulation of | 1.41 | preserved |
| 132 | regulation of | 1.37 | preserved |
| 133 | amide transp | 1.37 | preserved |
| 134 | organic subs | 1.56 | preserved |
| 135 | negative regu | 1.46 | preserved |
| 136 | catabolic pro | 1.59 | preserved |
| 137 | establishmer | 1.69 | preserved |
| 138 | protein locali | 1.42 | preserved |
| 139 | cellular respo | 14.96 | preserved |
| 140 | cellular prote | 2.14 | preserved |
| 141 | protein modi | 2.15 | preserved |
| 142 | cellular prote | 2.91 | preserved |
| 143 | macromolecu | 1.84 | preserved |
| 144 | protein meta | 2.12 | preserved |
| 145 | response to s | 11.04 | preserved |
| 146 | cellular macr | 2.26 | preserved |
| 147 | organonitrog | 1.88 | preserved |
| 148 | regulation of | 6.83 | preserved |
| 149 | organic subs | 1.43 | preserved |
| 150 | regulation of | 6.18 | preserved |
| 151 | biological reg | 5.26 | preserved |
| 152 | cellular proce | 2.14 | preserved |
| 153 | IRE1-mediate | 77.48 | perturbed |
| 154 | sequestering | 1.77 | perturbed |
| 155 | regulation of | 1.78 | perturbed |
| 156 | ATF6-mediat | 5.12 | perturbed |
| 157 | protein foldi | 4.99 | perturbed |
| 158 | endoplasmic | 84.06 | perturbed |
| 159 | integrated st | 4.77 | perturbed |
| 160 | PERK-mediat | 4.79 | perturbed |
| 161 | cellular respo | 81.21 | perturbed |
| 162 | cellular respo | 79.24 | perturbed |
| 163 | ER overload r | 2.74 | perturbed |
| 164 | positive regu | 2.75 | perturbed |
| 165 | response to u | 77.54 | perturbed |
| 166 | protein matu | 1.38 | perturbed |
| 167 | response to t | 75.8 | perturbed |
| 168 | regulation of | 1.33 | perturbed |
| 169 | ER-nucleus s | 8.47 | perturbed |
| 170 | response to e | 70.68 | perturbed |

| | | | |
|---|---|---|---|
| 171 | regulation of | 1.78 | perturbed |
| 172 | positive regu | 1.79 | perturbed |
| 173 | negative regi | 2.73 | perturbed |
| 174 | regulation of | 4.64 | perturbed |
| 175 | ubiquitin-dep | 2.15 | perturbed |
| 176 | positive regu | 1.33 | perturbed |
| 177 | ERAD pathwa | 1.89 | perturbed |
| 178 | maintenance | 1.81 | perturbed |
| 179 | protein foldir | 4.84 | perturbed |
| 180 | endoplasmic | 2.73 | perturbed |
| 181 | regulation of | 2.63 | perturbed |
| 182 | cellular respo | 2.53 | perturbed |
| 183 | cellular respo | 1.79 | perturbed |
| 184 | negative regi | 1.7 | perturbed |
| 185 | cellular respo | 37.64 | perturbed |
| 186 | regulation of | 2.7 | perturbed |
| 187 | proteasome- | 1.8 | perturbed |
| 188 | cellular respo | 34.63 | perturbed |
| 189 | proteasomal | 1.7 | perturbed |
| 190 | response to r | 1.59 | perturbed |
| 191 | Golgi vesicle | 1.48 | perturbed |
| 192 | cellular respo | 2.25 | perturbed |
| 193 | cellular respo | 30.84 | perturbed |
| 194 | response to c | 30.29 | perturbed |
| 195 | proteolysis ir | 2.03 | perturbed |
| 196 | ubiquitin-dep | 1.62 | perturbed |
| 197 | modification | 1.59 | perturbed |
| 198 | response to i | 1.58 | perturbed |
| 199 | cellular prote | 1.9 | perturbed |
| 200 | modification | 1.57 | perturbed |
| 201 | negative regi | 1.47 | perturbed |
| 202 | protein catal | 1.74 | perturbed |
| 203 | cellular respo | 1.39 | perturbed |
| 204 | response to s | 27 | perturbed |
| 205 | negative regi | 2.65 | perturbed |
| 206 | regulation of | 1.57 | perturbed |
| 207 | negative regi | 1.69 | perturbed |
| 208 | cellular homo | 1.67 | perturbed |
| 209 | negative regi | 1.65 | perturbed |
| 210 | response to c | 23.59 | perturbed |
| 211 | response to c | 1.44 | perturbed |
| 212 | signal transd | 21.6 | perturbed |
| 213 | cellular respo | 1.59 | perturbed |

**Table S4.** Gene coefficients for final pathway score models for four UPR pathways: XBP1s, PERK, ATF6, and RIDD. Coefficients are provided for each tumor type for which a model could be fitted.

| | BLCA | BRCA | CESC | CHOL | COAD | ESCA |
|---|---|---|---|---|---|---|
| ACADVL | 0 | 0 | 0 | -0.0266 | 0 | 0 |
| ADD1 | 0 | 0 | 0 | 0.0502 | 0 | 0 |
| ARFGAP1 | 0 | 0 | 0 | 0.0457 | 0 | 0 |
| ASNS | 0 | 0 | 0 | 0.064 | 0 | 0 |
| ATF3 | -5.25E-18 | -0.0055 | 0 | -0.0337 | 0 | 0 |
| ATF4 | 0 | 0 | 0 | 0.1517 | 0 | 0 |
| ATP6V0D1 | 0 | 0 | 0 | 0.0472 | 0 | 0 |
| BLOC1S1 | | | | | | |
| CALR | 0 | 0.0128 | 4.10E-18 | 0.0164 | 0 | 0 |
| CCL2 | 0 | 0 | 0 | -0.0014 | 0 | 0 |
| CD59 | | | | | | |
| CES1 | -6.55E-18 | 0 | 0 | -0.0808 | 0 | 0 |
| COL6A1 | | | | | | |
| CTDSP2 | 0 | 0 | 0 | 0 | 0 | 0 |
| CUL7 | 0 | 0 | 0 | 0.0585 | 0 | 0 |
| CXCL8 | 0 | 0 | 0 | 0.0214 | 0 | 0.0043 |
| CXXC1 | 0 | 0 | 0 | 0 | 0 | 0 |
| CYP2E1 | | | | | | |
| DCTN1 | 0 | 0 | 0 | 0.0774 | 0 | 0 |
| DDIT3 | 0 | 0 | 0 | 0.069 | 0 | 0 |
| DDX11 | 0 | 0 | 0 | 0.06 | 0 | 0.01 |
| DNAJB11 | 0 | 0 | 0 | 0.012 | 0 | 0 |
| DNAJB9 | 0 | 0 | 0 | 0 | 0 | 0 |
| DNAJC3 | 0 | 0 | 0 | 0 | 0 | 0 |
| EDEM1 | 0 | 0 | 0 | 0 | 0 | 0 |
| ERN1 | | | | | | |
| EXTL1 | 0 | 0 | 0 | 0 | 0 | 0 |
| EXTL2 | 0 | 0 | 0 | 0 | 0 | 0 |
| EXTL3 | 0 | 0 | 0 | 0 | 0 | 0 |
| FKBP14 | 0 | 0 | 0 | 0.0043 | 0 | 0 |
| GALNT2 | | | | | | |
| GAS2 | 0 | 0 | | | | 0 |
| GEMIN5 | | | | | | |
| GFPT1 | 0 | 0 | 0 | 0.0161 | 0 | 0 |
| GOSR2 | 0 | 0 | 0 | 0 | 0 | 0 |
| GPC3 | 0 | -0.0054 | | | | 0 |
| GSK3A | 0 | 0 | 0 | 0.0791 | 0 | 0 |
| HDGF | 0 | 0.0124 | 0 | 0 | 0 | 0.034 |
| HERPUD1 | 0 | 0 | 0 | 0.0968 | 0 | 0 |
| HGSNAT | | | | | | |
| HSP90B1 | 0 | 0 | 0 | 0.0953 | 3.01E-17 | 0 |
| HSPA5 | 0 | 0 | 0 | 0.0612 | 0 | 0 |

**Table S5.** Regression model coefficients, p values and 95% confidence intervals for a pancancer OLS model linking SCNA and four UPR pathway activity scores to CYT levels, including tumor type as a covariate, but excluding IHC level.

| | Coeff | P value | 95% CI |
|---|---|---|---|
| SCNA | -0.300 | 1.42e-119 | -0.325, -0.275 |
| XBP1S | 0.092 | 0.062 | -0.005, 0.188 |
| PERK | -0.376 | 8.61e-13 | -0.478, -0.273 |
| ATF6 | 0.201 | 0.001 | 0.078, 0.324 |
| RIDD | -0.142 | 0.010 | -0.249, -0.035 |
